# FAK family proteins regulate *in vivo* breast cancer metastasis *via* distinct mechanisms

**DOI:** 10.1101/2023.10.27.564212

**Authors:** Alessandro Genna, Joel Alter, Martina Poletti, Tomer Meirson, Tal Sneh, Michal Gendler, Natalia Saleev, George S. Karagiannis, Yarong Wang, Dianne Cox, David Entenberg, Maja H. Oktay, Tamas Korcsmaros, John S. Condeelis, Hava Gil-Henn

## Abstract

Breast cancer is the most commonly diagnosed malignancy and the major leading cause of tumor-related deaths in women. It is estimated that the majority of breast tumor-related deaths are a consequence of metastasis, to which no cure exists at present. The FAK family proteins Proline-rich tyrosine kinase (PYK2) and focal adhesion kinase (FAK) are highly expressed in breast cancer, but the exact cellular and signaling mechanisms by which they regulate *in vivo* tumor cell invasiveness and consequent metastatic dissemination are mostly unknown. Using a PYK2 and FAK knockdown xenograft model we show here, for the first time, that ablation of either PYK2 or FAK decreases primary tumor size and significantly reduces Tumor MicroEnvironment of Metastasis (TMEM) doorway activation, leading to decreased intravasation and reduced spontaneous lung metastasis. Intravital imaging analysis further demonstrates that PYK2, but not FAK, regulates a motility phenotype switch between focal adhesion-mediated fast motility and invadopodia-dependent, ECM-degradation associated slow motility within the primary tumor. Furthermore, we validate our *in vivo* and intravital imaging results with integrated transcriptomic and proteomic data analysis from xenograft knockdown tumors and reveal new and distinct pathways by which these two homologous kinases regulate breast tumor cell invasiveness and consequent metastatic dissemination. Our findings identify PYK2 and FAK as novel mediators of mammary tumor progression and metastasis and as candidate therapeutic targets for breast cancer metastasis.

## INTRODUCTION

Breast cancer is the most commonly diagnosed malignancy and the major leading cause of tumor-related deaths among women worldwide^1,2^. The different subtypes of breast tumors are associated with diverse clinical outcomes and therapeutic approaches^3^. Hormonal therapies are a first line treatment in estrogen receptor (ER)- and progesterone receptor (PR)-positive breast tumors^4^, while anti-HER2 antibodies and tyrosine kinase inhibitors are used to treat HER2-positive breast cancer^5^. Unfortunately, no targeted therapies besides chemotherapy have been approved for triple negative breast cancer (TNBC), to date^6^. Despite recent advances in detection and therapeutics of breast cancer, chemotherapy resistance, disease relapse, and metastasis remain as major challenges in the treatment of breast cancer patients at present^7,8^. It has been estimated that a majority of deaths due to breast cancer are a consequence of metastasis, the spread of tumor cells to other tissues and organs in the body and the formation of secondary tumors which impair the function of vital organs such as lungs, liver, and brain. While researchers have identified therapeutic strategies to shrink the primary tumor or slow its growth, no treatment to permanently eradicate metastasis exists at present^9^.

Focal adhesion kinase (FAK) and proline-rich tyrosine kinase 2 (PYK2) define a family of non-receptor tyrosine kinases that share high amino acid identity, common phosphorylation sites, and a similar domain structure, which includes N-terminal 4.1, ezrin, radixin, moesin (FERM) domain; a tyrosine kinase domain; proline-rich regions; and a C-terminal focal adhesion targeting (FAT) domain. While FAK is ubiquitously expressed, PYK2 expression is more restricted and is especially high in the central nervous system and in hematopoietic cells^10^. Several studies reported overexpression of FAK in breast, colon, ovary, pancreas, and prostate cancers, as well as in other cancers^11,12^. FAK is involved in several different cellular processes that contribute to tumor initiation, progression, and metastasis in breast cancer, such as proliferation, cell cycle progression^13^, survival, epithelial-to-mesenchymal transition (EMT), breast cancer stem cell (CSC) self-renewal and maintenance, tumor cell motility, mechanotransduction, and regulation of the breast tumor microenvironment (TME)^14^. Overexpression of FAK in human breast cancer correlates with more aggressive and invasive breast carcinoma^12^ and with significantly shorter metastasis-free survival^15^. The role of FAK in mammary tumor progression and metastasis has previously been demonstrated in mouse models using FAK specific deletion in the mammary epithelium^15–18^, however the *in vivo* cellular and signaling mechanisms by which FAK regulates tumor cell motility and invasiveness have not been thoroughly explored to date.

Similar to FAK, high expression of PYK2 has been observed in several human tumors including glioblastoma, hepatocellular carcinoma, non-small cell lung carcinoma, prostate cancer, as well as early and advanced breast cancer^19^. PYK2 regulates several cellular processes which contribute to tumorigenesis and invasiveness of breast cancer cells, such as tumor cell growth, cell motility and invasion, EMT and stemness, and in regulation of tumor cell-macrophage communication that leads to macrophage infiltration and consequent tumor growth and angiogenesis^20^. PYK2 expression is significantly up-regulated in recurrent human breast cancer^21^, and its high expression correlates with higher tumor grade and with lymph node invasion^22^. A role for PYK2 in regulating primary tumor growth and metastatic outgrowth has been demonstrated using mouse models^21,23^, but whether and how PYK2 is involved in the *in vivo* invasiveness and metastatic dissemination of breast cancer cells, and does it do so *via* molecular and cellular mechanisms similar or distinct from the closely related FAK, is unknown at present.

Within primary breast tumors, transient vascular opening and its associated tumor cell intravasation, occur simultaneously and exclusively at microanatomical structures called Tumor MicroEnvironment of Metastasis (TMEM) ^24,25^. Each functional TMEM doorway is composed of three different cell types in direct and stable physical contact: a tumor cell highly expressing the actin regulatory protein Mammalian enabled (MENA), a perivascular Tie2^hi^/Vegf^hi^ macrophage, and an Ang2 secreting endothelial cell^24,26^. TMEM doorways have been identified in mouse and human mammary carcinomas, and their density (TMEM doorway score) correlates with metastatic outcome in breast cancer patients^27–31^. Metastatic cancer cells intravasate into blood vessels and disseminate in the body *via* TMEM-associated transient vascular openings (TAVO)^32–34^. Trans-endothelial migration of invasive tumor cells through TMEM doorways requires penetrating the endothelial basement membrane, which is achieved by using F-actin rich protrusive membrane structures with ECM degrading activity called invadopodia^32,35,36^. Invadopodia were identified in several invasive cancer cell lines including breast cancer^37^ as well as in primary tumor cells obtained from patients^38,39^. Previous publications demonstrate a direct molecular link between invadopodia assembly and function and metastasis in mouse models^40,41^. Importantly, the existence of ECM-degrading invadopodia was previously demonstrated in invasive breast tumor cells to be involved in transendothelial migration by intravital microscopy in mice^34,42^.

We have previously shown that PYK2 regulates invadopodia maturation and activation by mediating epidermal growth factor (EGF)-induced tyrosine phosphorylation of cortactin, both directly, and indirectly *via* Src-mediated Arg activation. Both phosphorylation events lead to actin polymerization in invadopodia, ECM degradation, and tumor cell invasiveness^43^. While FAK does not localize to invadopodia, it coordinates, together with PYK2, the balance between focal adhesion-mediated motility and invadopodia-mediated invasiveness by controlling Src kinase localization. This coordination between PYK2-mediated invasion and FAK-mediated migration is necessary for breast cancer cell invasiveness^43,44^. Whether and how PYK2 and FAK use these distinct cellular mechanisms for regulating *in vivo* invasion and metastatic dissemination is unknown at present.

In this study, we elucidate the cellular and signaling mechanisms by which PYK2 and FAK regulate *in vivo* invasiveness and metastasis using a xenograft mouse model of breast cancer. Our results show that, knockdown of either PYK2 or FAK significantly reduce *in vivo* invasiveness, the number of circulating tumor cells, and spontaneous lung metastasis. We find this reduced metastasis to correlate with decreased TMEM doorway-dependent vascular opening, but not with TMEM doorway assembly. Intravital imaging experiments further demonstrate that PYK2, but not FAK, regulates the switch between focal adhesion-mediated fast motility and invadopodia-dependent ECM degradation-associated slow motility within the primary tumor. Thus, although highly homologous, PYK2 and FAK appear to regulate metastasis *via* distinct mechanisms. Integrated transcriptomics, proteomics, and protein-protein interaction data from primary knockdown tumors further support our *in vivo* and intravital findings while revealing new pathways by which PYK2 and FAK regulate tumorigenesis and metastasis. Collectively, our findings elucidate the cellular and signaling mechanisms by which PYK2 and FAK regulate breast cancer metastasis and identifies these kinases as potential therapeutic targets in breast cancer.

## RESULTS

### FAK and PYK2 act synergistically to promote breast cancer metastasis

We previously suggested a model in which PYK2 and FAK modulate cancer cell invasion by coordinating the balance between focal adhesion-mediated migration and invadopodia-dependent ECM invasion^43,44^. According to this model, both Pyk2-mediated invasion and FAK-mediated migration are necessary for breast cancer cell invasiveness and subsequent metastatic dissemination^43,44^. To characterize the involvement of PYK2 and FAK in breast cancer metastasis in patients, we integrated DNA, RNA, and protein expression data of breast cancer tissues from The Cancer Genome Atlas (TCGA) and stratified them by hormone- and HER2-receptor status (**Figure 1A**). A comparison of copy number alterations (CNA) (**Figure 1B**), mRNA expression (**Figure 1C**), and protein expression (**Figure 1D**) demonstrated similar distribution among all hormone- and HER2-receptor statuses. These data indicate that PYK2 (*PTK2B*) and FAK (*PTK2*) expression profile is similar across all receptor statuses and is not unique to any of the four primary molecular subtypes in breast cancer. To assess the involvement of PYK2 and FAK in metastatic dissemination and the clinical potential of their inhibition, we analyzed microarray expression levels of 1,650 breast cancer samples from TCGA. The survival analysis showed a significant association between poor distant metastasis-free survival (DMFS) and high mRNA levels of *PTK2* (HR 1.49, 95% CI 1.21-1.82; *p* = 1.6e^-4^) (**Figure 1E**) but not with *PTK2B* (HR 1.06, 95% CI 0.86-1.30; *p* = 0.62) (**Figure 1F**). However, the combined effect of *PTK2* and *PTK2B* overexpression considerably reduced DMFS estimates (HR 1.68, 95% CI 1.22-2.30; *p* = 1.9e^-3^) with a clinically significant synergy index (SI) of 1.24 ^45^ (**Figure 1G**), indicating that PYK2 and FAK act synergistically to promote breast cancer metastasis.

**Figure 1.**
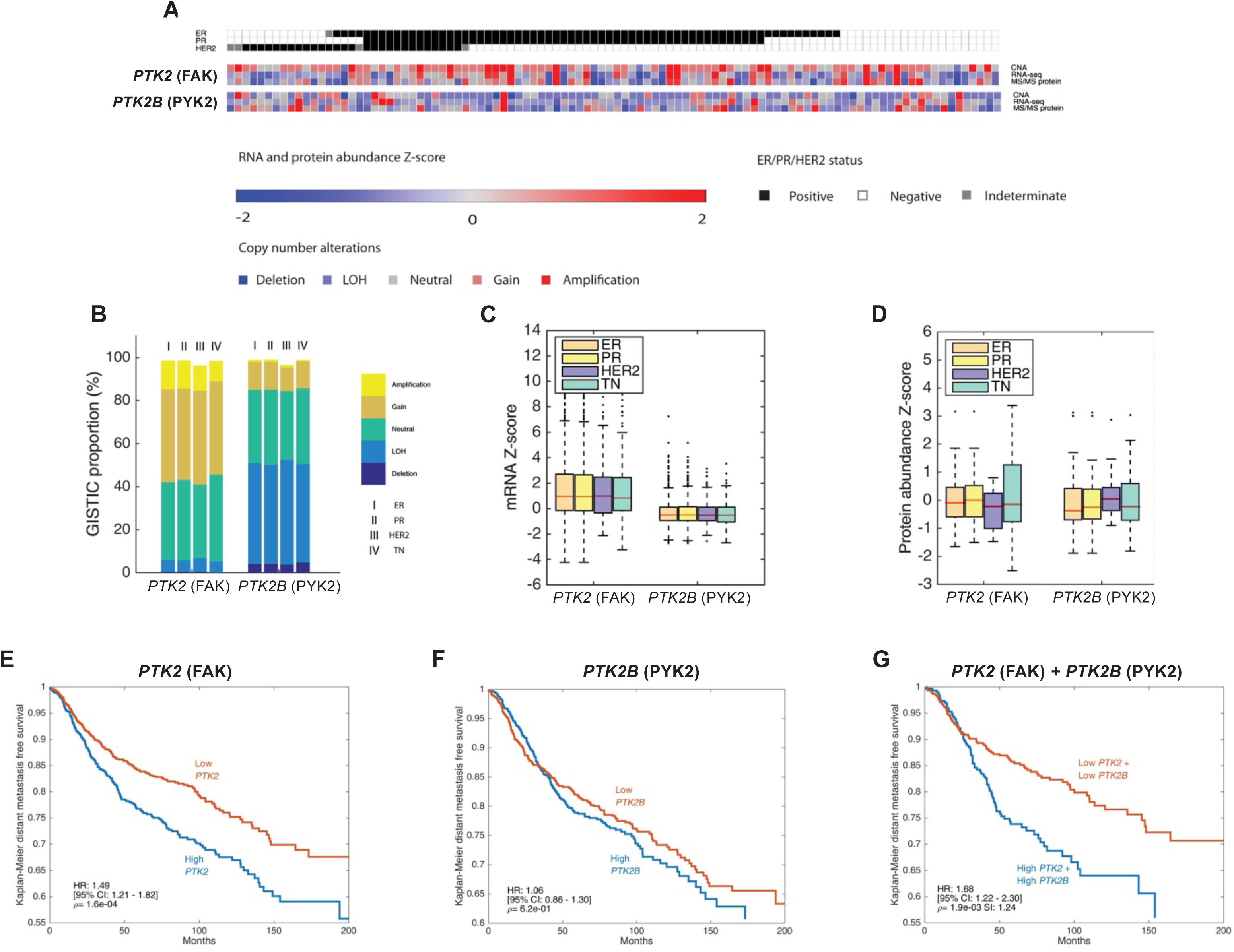
PYK2 and FAK act synergistically to promote breast cancer metastasis. **(A)** Heat map of copy number alteration (CNA), mRNA expression, and protein expression of FAK (*PTK2*) and PYK2 (*PTK2B*) across 102 tumor samples based on the CPTAC database. ER, estrogen receptor positive; PR, progesterone receptor positive; HER2, HER2 positive; TN, triple negative. GISTIC, mRNA and protein abundance Z-scores are shown for each sample. **(B)** GISTIC analysis of 1,905 samples originated from TCGA with different hormone receptor statuses. **(C)** mRNA expression analysis of 1,905 samples originated from TCGA with different hormone receptor statuses. **(D)** Comparison of protein abundance in 102 samples from different hormone receptor statuses. The boxes in C, D are delimited by the lower and upper quartile, the horizontal red line indicates the sample median, the notches represent the 95% confidence interval (CI), and whiskers extend to the most extreme point which is no more than 1.5 times the interquartile range from the box. Outliers are shown as points beyond boxplot whiskers. P values were calculated using two-tailed Student’s *t*-test. **(E-G)** Kaplan-Meier curves of DMFS in 1,650 breast cancer cases. Microarray data (Affymatrix) were obtained from the NCBI Gene Expression Omnibus (GEO) data repository. Tumor samples were split into high and low expression groups based on mRNA gene expression with z-score cut-off value of 0. Shown are DMFS survival curves for *PTK2* (E), *PTK2B* (F), and their combination (G). P values were calculated by log-rank test, hazard ratio (HR) and 95% CI are shown. Synergy index (SI) is indicated for evaluation of *PTK2+PTK2B* in (G).

### Knockdown of PYK2 or FAK decreases primary tumor size

For this study, we used the MDA-MB-231 cell line, an established human triple-negative, basal-like breast adenocarcinoma cell line that is widely used for metastasis studies due to its ability to grow orthotopic tumors that spontaneously metastasize to the lungs in mice. Using this cell line, we generated stable knockdown of *PTK2B* and *PTK2* in MDA-MB-231 with two distinct short-hairpin RNA (shRNA) sequences for each (PYK2-KD1, PYK2-KD2 and FAK-KD1, FAK-KD2 cell lines) or an empty vector as control. As shown in **Figure 2A**, all four sequences resulted in significant knockdown of either PYK2 or FAK expression.

**Figure 2.**
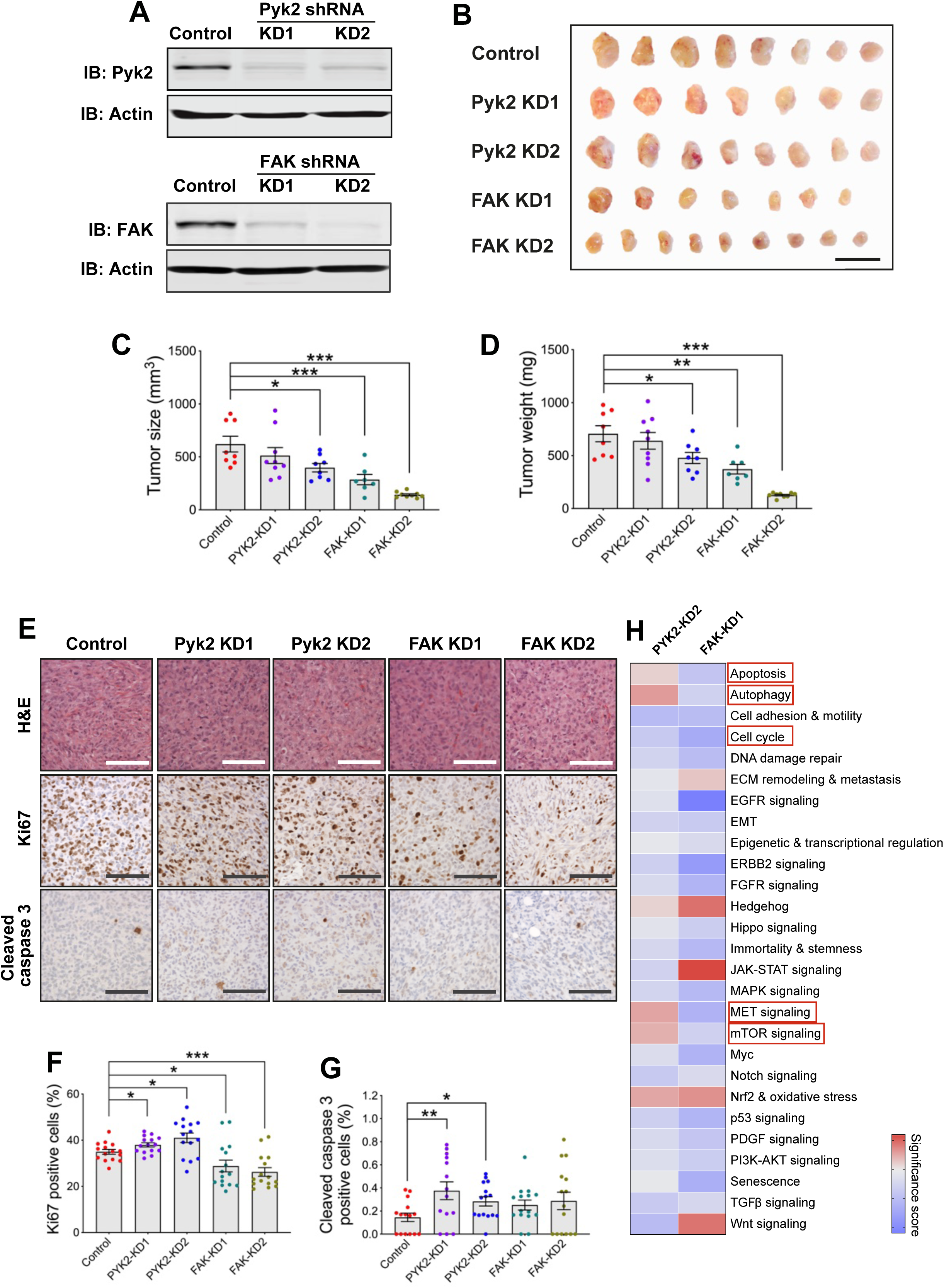
Knockdown of PYK2 or FAK significantly decreases primary tumor size. **(A)** Immunoblots showing knockdown of PYK2 or FAK in MDA-MB-231/Dendra2 cell lines by two different shRNA sequences for each gene, PYK2-KD1, PYK2-KD2 and FAK-KD1, FAK-KD2, relative to control empty vector expressing cells. Actin is used as a loading control. **(B)** Orthotopic tumors were generated either by injection of control, PYK2-KD, or FAK-KD MDA-MB-231/Dendra2 cells into the mammary gland of SCID mice. Ten weeks following injection, mice were sacrificed, and tumors were isolated and measured. Shown are representative images of tumors from control, PYK2-KD, and FAK-KD cells. Scale bar, 2 cm. **(C-D)** Quantification of primary tumor size (C) and weight (D) of orthotopic tumors generated by injection of control, PYK2-KD, or FAK-KD cells. *n* = 7-11 mice per group. The experiment was repeated twice with similar results. **(E-G)** Representative images (E) and quantification of proliferating cells (Ki67; F) and apoptotic cells (cleaved caspase 3; G). *n* = 15 random fields from 3 tumors per condition. Scale bar, 50 μm. **(H)** Enrichment analysis of transcriptome in PYK2-KD2 and FAK-KD1 tumors, that was measured by NanoString. All tumors were collected 10 weeks following mammary gland injection. *n* = 5 tumors per group.

To test how the loss of PYK2 or FAK affects cancer progression *in vivo*, we injected the stable cell lines into the mammary gland of severe combined immunodeficiency (SCID) mice. At 10 weeks following injection, primary tumors were isolated and tumor size and weight were measured. As demonstrated in **Figure 2B-D**, tumors arising from FAK-KD1 and FAK-KD2 and tumors arising from PYK2-KD2 were significantly smaller than tumors generated from the control cell line. To gain insight into the mechanisms responsible for the smaller size of both FAK-KD and PYK2-KD tumors, we used histological markers for proliferation and apoptosis, processes that may affect primary tumor growth. As demonstrated in **Figure 2E, F**, PYK2-KD tumors showed increased proliferation while significantly decreased proliferation was observed in FAK-KD tumors. Moreover, PYK2-KD tumors showed increased apoptosis marker staining (**Figure 2E, G**).

To further elucidate the molecular and signaling mechanisms leading to smaller tumor size in PYK2 and FAK knockdown, we examined gene expression in the knockdown tumors relative to control using NanoString (**Figure 2H**). Interestingly, while the transcription of cell cycle associated genes was significantly down-regulated in both PYK2-KD and FAK-KD tumors (pathway significance score: −1.421 (Pyk2-KD2), −2.682 (FAK-KD1)) (**Supplementary** Figure 1A), a significant increase in mTOR signaling (pathway significance score: 0.941) (**Supplementary** Figure 2B) and in MET signaling (pathway significance score: 1.115) (**Supplementary** Figure 2C) was observed in PYK2-KD tumors, which may explain their higher proliferation rate, while the transcription of apoptosis- and autophagy-associated genes was significantly up-regulated in these tumors (pathway significance score: 0.43 (apoptosis), 1.248 (autophagy)), which could explain their smaller size (**Supplementary** Figure 2D**, E**). Together, these data imply that while knockdown of FAK results in smaller tumors due to decreased proliferation, the decreased size of PYK2-KD tumors results from enhanced apoptosis.

### Pyk2 or FAK knockdown significantly reduces *in vivo* invasiveness, intravasation, and lung metastasis in xenograft mice

We next sought to determine whether PYK2 or FAK control the invasiveness of tumor cells *in vivo*, within the context of the primary tumor. For this purpose, we used the *in vivo* invasion assay in which tumor cells can be collected from equal sized MDA-MB-231 primary tumors in response to an EGF gradient^46^ (**Figure 3A**). The total number of tumor cells invading towards EGF in PYK2-KD and FAK-KD tumors was significantly reduced compared with control tumors of the same size (**Figure 3B**). Given the reduced invasiveness of PYK2-KD and FAK-KD tumor cells, we next tested how subsequent steps in metastasis were affected. Using an established circulating tumor cell (CTC) assay^47^ (**Figure 3C**), we found that both PYK2-KD and FAK-KD tumor-bearing mice had significantly fewer CTCs in their blood as compared with mice bearing control tumors of similar size (**Figure 3D**). This finding indicates that both PYK2 and FAK are necessary for intravasation of tumor cells into the bloodstream.

**Figure 3.**
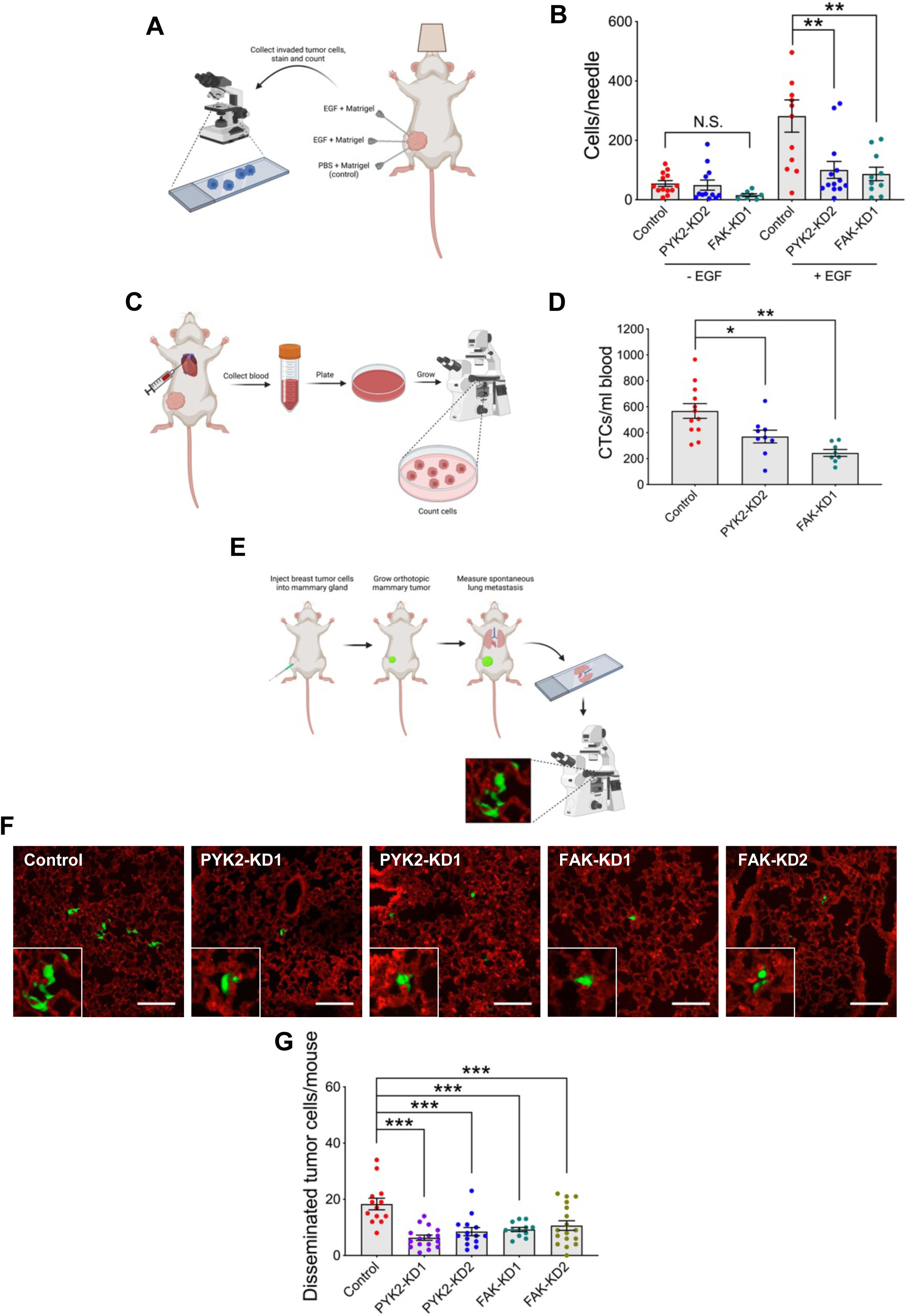
Knockdown of either PYK2 or FAK significantly reduces *in vivo* invasion, intravasation, and lung metastasis in xenograft mice. **(A-B)** *In vivo* invasion towards EGF was measured in orthotopic primary tumors generated with either control, PYK2-KD2, or FAK-KD1 MDA-MB-231/Dendra2 cells. Total cells were counted by DAPI staining. *n* = 5-8 mice per group. **(C-D)** Assay cartoon (C) and quantification (D) of circulating tumor cells in mice bearing control, PYK2-KD2, or FAK-KD1 mammary tumors. *n* = 12 (control), *n* = 9 (PYK2-KD), *n* = 8 (FAK-KD) mice. **(E-G)** Spontaneous metastasis cartoon (E) and representative images (F) of lung sections from mice bearing orthotopic tumors generated by control, PYK2-KD and FAK-KD MDA-MB-231/Dendra2 cells. Green, tumor cells; red, lung blood vessels. Scale bar, 100 μm. **(G)** Quantification of single disseminated tumor cells. *n* = 14 (control), *n* = 16 (PYK2-KD1), *n* = 14 (PYK2-KD2), *n* = 12 (FAK-KD1), *n* = 17 (FAK-KD2) mice from two independent experiments.

At the final stage of metastasis, cancer cells extravasate into the target organ to establish new metastatic colonies. Because the knockdown of either PYK2 or FAK significantly decreased *in vivo* invasion and the number of CTCs that originated from the primary knockdown tumors, we hypothesized that metastatic dissemination to the lungs of tumor bearing mice would also be decreased. As demonstrated in **Figure 3E-G**, mice bearing PYK2-KD or FAK-KD orthotopic tumors exhibited significantly fewer disseminated tumor cells than control mice bearing equal size tumors. Collectively, these data suggest that both PYK2 and FAK have a crucial role in breast cancer metastatic dissemination by promoting invasiveness and intravasation of cancer cells from the primary tumor.

### TMEM doorway-dependent vascular opening, but not TMEM doorway assembly, is significantly reduced in PYK2 and FAK knockdown tumors

The significant decrease in *in vivo* invasion, CTCs, the number of disseminated tumor cells in lung, and lung metastases in both PYK2-KD and FAK-KD tumor bearing mice (**Figure 3**) led us to examine whether knockdown of either kinase affects TMEM doorway assembly or activation. Control and knockdown tumors were sectioned and subjected to triple TMEM doorway staining and analysis. Interestingly, TMEM doorway score was not affected by knockdown of either PYK2 or FAK (**Figure 4A, B, C**). Next, we sought to determine whether TMEM doorway functionality is altered by PYK2 or FAK knockdown. TMEM doorway functionality can be assessed by introducing into the blood serum a high-molecular weight fluorescently labeled dextran which is incapable of crossing the endothelial barrier except through TMEM doorways during TAVO, and then measuring the levels of extravascular dextran in the perivascular region around TMEM doorways^48^. A fundamental event for tumor cell intravasation through TMEM doorways is the transient opening of the blood vessel at the TMEM doorway which allows the tumor cell to enter the bloodstream^24^. Indeed, reduced levels of extravascular dextran were observed in PYK2-KD and FAK-KD tumors comparing to control tumors, indicating reduced TAVO (**Figure 4D, E**).

**Figure 4.**
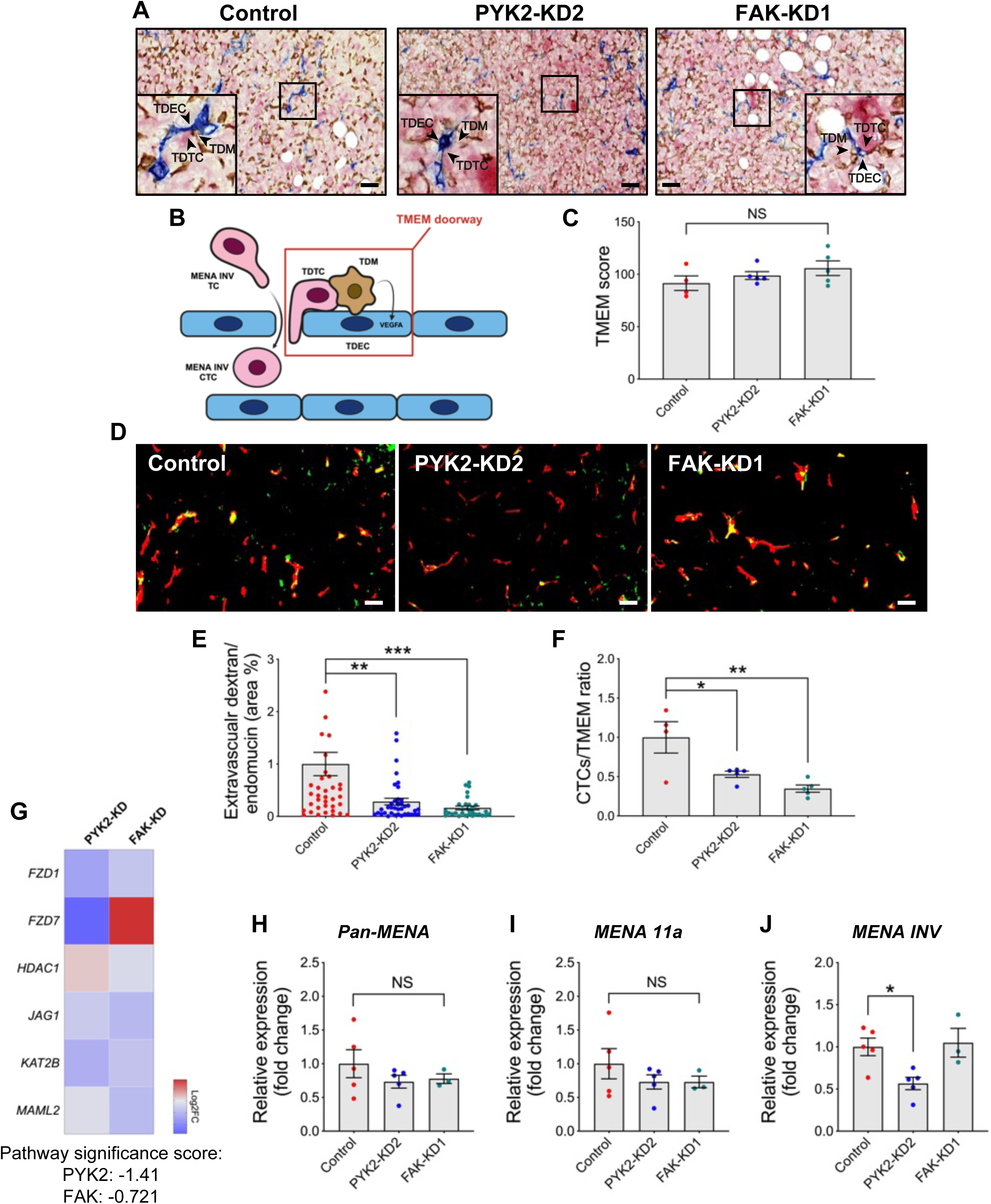
TMEM doorway associated vascular opening, but not TMEM-doorway assembly, is significantly reduced in PYK2 and FAK knockdown tumors. **(A)** Representative images of IHC triple staining of TMEM doorways in control, PYK2-KD2, and FAK-KD1 tumors. Inset, TMEM doorway composed of a tumor cell (pink), endothelial cell (blue) and a macrophage (brown). MENA^INV^ TC, tumor cell expressing the invasive isoform of MENA; TDTC, TMEM doorway tumor cell; TDM, TMEM doorway macrophage; MENA^INV^ CTC, circulating tumor cell expressing the invasive isoform of MENA. Scale bar, 50 μm. **(B)** Cartoon illustration of a TMEM doorway, composed of a TMEM doorway tumor cell (TDTC), a TMEM doorway macrophage (TDM), and a TMEM doorway endothelial cell (TDEC), all in direct and stable contact. VEGFA secretion from the TDM leads to opening of the TMEM doorway. MENA^INV^ expression in tumor cells near TMEM doorways (MENA^INV^ TC) enables these cells to migrate through the open doorway and become a circulating tumor cell (MENA^INV^ CTC). **(C)** TMEM doorway score of control, PYK2-KD2, and FAK-KD1 tumors. **(D)** Representative blood vessel (red) and extravascular dextran (green) masks, as obtained by immunofluorescence of control, PYK2-KD, and FAK-KD tumors, showing TMEM doorway-associated vascular opening (yellow). Scale bar, 50 μm. **(E)** Quantification of extravascular dextran area normalized to blood vessel area in control and KD tumors. *n* > 40 fields from each tumor group. **(F)** Ratio of CTCs to TMEM doorways. *n* = 4 tumors (control), *n* = 5 tumors (PYK2-KD2, FAK-KD1). **(G)** Expression of genes involved in the NOTCH1 pathway in PYK2-KD and FAK-KD tumors, as detected by NanoString analysis. Significance score of the pathway in tumors: −1.41 (PYK2-KD2), −0.721 (FAK-KD1). **(H-J)** qPCR expression analysis of *pan-MENA* (H), *MENA11a* (I), and *MENA^INV^* in control, PYK2-KD2, and FAK-KD1 tumors.

Moreover, the ratio of CTCs to TMEM doorway score was significantly reduced in PYK2-KD and FAK-KD tumors (**Figure 4F**). Assuming that TMEM doorway activity has a constant rate and that the intravasation of invasive cancer cells into blood vessels is performed *via* TMEM doorways only (Harney et al 2015), these results indicate that reduced TMEM doorway functionality leads to decreased intravasation in the knockdown tumors. Together, these data indicate that lower metastatic dissemination in PYK2 and FAK knockdown tumor bearing mice results from lower TMEM doorway-associated intravasation, but not TMEM doorway score.

It was previously demonstrated that macrophage-tumor cell contact enhances NOTCH1 signaling and stimulates up-regulation of MENA^INV^ expression in the contacted tumor cell, triggering the assembly and activation of invadopodia and subsequent trans-endothelial migration and tumor cell dissemination at TMEM doorways^34,38,49,50^. To examine whether knockdown of PYK2 or FAK correlates with changes in NOTCH1 signaling, we used NanoString analysis for expression of genes in the NOTCH1 pathway in PYK2-KD and FAK-KD tumors relative to control tumors. Importantly, both tumors lacking PYK2 and tumors lacking FAK showed reduced expression of genes involved in NOTCH1-mediated signaling (pathway significance score: −1.41 (PYK2-KD), −0.721 (FAK-KD)) (**Figure 4G**). Additionally, while no changes in expression of *pan-MENA* or the *MENA11a* isoform were observed in the PYK2-KD or FAK-KD tumors (**Figure 4H, I**), a significant decrease in *MENA^INV^*expression was observed in PYK2-KD tumors, but not in FAK-KD tumors (**Figure 4J**). All in all, these findings suggest that both PYK2 and FAK regulate NOTCH1 signaling within tumor cells, leading to TMEM doorway activation and subsequent tumor cell intravasation.

### PYK2, but not FAK, localizes to invadopodia-like structures *in vivo*

PYK2 has previously been shown to localize to cortactin- and TKS5-containing, matrix degrading invadopodia *in vitro*^43^ and our data presented here suggest it is involved in controlling breast cancer metastasis *in vivo*. However, the association of PYK2 with invadopodia-like tumor cellular protrusions has never been examined *in vivo*. To investigate the cellular localization of PYK2 and FAK within the primary tumor, we examined tissue sections of orthotopic xenograft tumors that were generated from MDA-MB-231 cells expressing PYK2-GFP or FAK-GFP, cortactin-TagRFP as an invadopodia marker, and MMP Sense, a protease-activatable fluorescent *in vivo* imaging agent that is optically silent upon injection, but which produces a fluorescent signal following cleavage by MMPs. As demonstrated in **Figure 5A-C**, co-localization of PYK2, cortactin, and fluorescently activated MMP Sense was observed in primary tumor sections, supporting enrichment of PYK2 in matrix-degrading tumor cell protrusions. Interestingly, and in line with previous observations, no localization of FAK to invadopodia-like structures was observed in primary tumors. These data validate our previous *in vitro* findings demonstrating that PYK2, but not FAK, localizes to invadopodia^43^.

**Figure 5.**
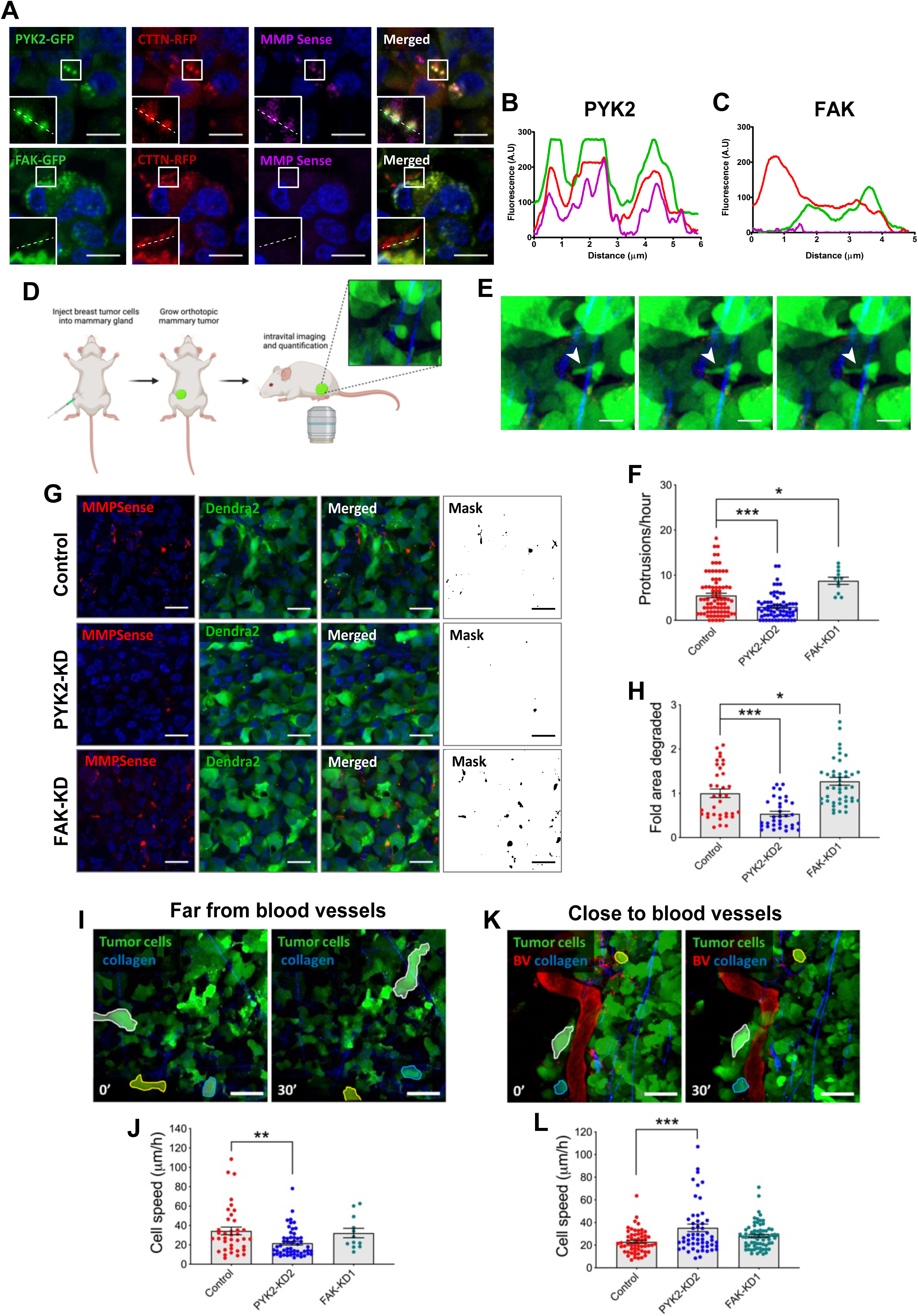
Knockdown of PYK2 decreases, while knockdown of FAK increases, protrusion frequency and *in vivo* ECM degradation. **(A)** Localization of PYK2 or FAK was visualized using MMP Sense (magenta) in orthotopic xenograft tumors generated from MDA-MB-231 cells stably expressing PYK2-GFP or FAK-GFP and cortactin-TagRFP. Co-localization of PYK2, but not FAK, with activated MMP Sense can be seen. Scale bar, 10 μm. **(B, C)** Intensity profiles of PYK2/FAK (green line), cortactin (red line), and MMP Sense (magenta line), illustrating enrichment of PYK2 (B), but not FAK (C), in matrix-degrading protrusions. **(D)** Cartoon description of imaging and quantification of invadopodia-like protrusions by intravital imaging. **(E)** Representative intravital images of a fluorescent tumor cell (dendra2, green) sending a protrusion (white arrowhead) towards a collagen fiber (second harmonic generation, blue) inside a mammary tumor. Images were extracted from Supplementary Video 1. Scale bar, 25 μm. **(F)** Quantification of protrusions in control, PYK2-KD, and FAK-KD tumors in regions close to blood vessels (<100 μm from a blood vessel). **(G)** Representative images (top panels) and quantification masks (bottom panel) of MMP activity in orthotopic xenograft tumors generated by control, PYK2-KD, or FAK-KD MDA-MB-231/Dendra2 cells (green). ECM degradation was visualized using fluorescently activated MMP Sense (red), and DAPI (blue). Scale bar, 25 μm. **(H)** Quantification of ECM degradation in tumors. *n* = 35 (control), *n* = 36 (PYK2-KD), *n* = 45 (FAK-KD) fields from three mice in each group. **(I, K)** Representative multi-photon images of MDA-MB-231/Dendra2 motile cells (green) migrating in orthotopic xenograft tumors in a region far from a blood vessel (I) and in a region close to blood vessel (K). Blood vessels are labeled by rhodamine-conjugated dextran (red) and collagen fibers are imaged by second harmonic generation (blue). Scale bar, 25 μm. Images were extracted from Supplementary Videos 2, 4. **(J, L)** Quantification of cell speed in control, PYK2-KD, and FAK-KD tumors in regions far from blood vessels (>100 μm from a blood vessel; B) and close to blood vessels (<100 μm from a blood vessel; D). *n* = 18-71 cells from 3-5 mice in each group.

### Knockdown of PYK2 decreases, while knockdown of FAK increases, protrusion frequency and *in vivo* ECM degradation

To elucidate single tumor cell behavior within the primary tumor of live mice, we used high-resolution intravital imaging (IVI) of control and knockdown MDA-MB-231 cells expressing Dendra2. Using this same method, it was previously shown that within a primary xenograft tumor, single tumor cells send small protrusions that have been identified as invadopodia based on morphological, molecular, and functional assays, and are associated with ECM degradation^34,42^. Examination of tumor cell protrusions in control and knockdown tumors revealed a significant decrease in protrusion frequency in PYK2-KD tumors and a significant increase in FAK-KD tumors (**Figure 5D, E, F and Supplementary Video 1**).

Our previous *in vitro* data demonstrated that knockdown of PYK2 decreases, while knockdown of FAK increases, MMP-mediated matrix degradation by invadopodia and subsequent breast tumor cell invasiveness^43^. To gain further insight into the *in vivo* invasion mechanism, and to examine whether knockdown of PYK2 or FAK impedes MMP-mediated matrix degradation within the primary tumor, we performed the *in vivo* MMP activity assay. Mice bearing fluorescently labeled control, PYK2-KD, or FAK-KD tumors were injected with MMP Sense before sacrifice. Tumors were then sliced and imaged by confocal microscopy and the fluorescent signal that was generated by MMP-mediated activity within the primary tumor was quantified. As demonstrated in **Figure 5G-H**, the fluorescence signal that is directly a result of MMP activation and ECM degradation was significantly decreased in PYK2-KD tumors and significantly increased in FAK-KD, compared to control tumors.

Collectively, these data support our previous *in vitro* findings^43^ indicating that PYK2 and FAK oppositely regulate the assembly of invadopodium precursors in breast cancer cells. Accordingly, decreased tumor cell invasion, despite elevated invadopodium precursor assembly and MMP-mediated ECM degradation in FAK-depleted cells was previously explained by re-localization of Src kinase to invadopodium precursors and subsequent formation and activation of these structures^43,44^.

### PYK2 regulates a motility phenotype switch in primary breast tumors

Invasive breast tumor cells show two motility phenotypes within primary tumors: focal adhesion-mediated fast motility on ECM fibers, and slow invadopodia-mediated ECM degradation-associated motility. Both of these motility phenotypes contribute to metastasis; tumor cells use fast locomotion to translocate to perivascular regions where they stop due to ECM composition and stiffness and switch to an invadopodium-associated phenotype that results in their intravasation and dissemination^34,42^. The involvement of PYK2 and FAK in regulation of focal adhesion-mediated migration and invadopodia-mediated tumor cell invasiveness has led us to hypothesize that one or both kinases may regulate the switch between the two motility phenotypes within the primary tumor.

To test this hypothesis, we measured the motility of individual tumor cells in control, PYK2-KD, and FAK-KD tumors in regions of the tumor that are far (>100 μm) from blood vessels, where fast locomotion occurs, and in regions close (<100 μm) to blood vessels, where slow locomotion occurs^42^. Interestingly, in regions far from blood vessels, the speed of PYK2-KD tumor cells was significantly reduced compared to control cells, while no significant change was observed in the speed of FAK-KD cells (**Figure 5I, J and Supplementary Videos 2, 4**). In regions close to blood vessels, the speed of PYK2-KD tumor cells was significantly increased while no significant change in the speed of FAK-KD cells was observed (**Figure 5K, L and Supplementary Videos 3, 5**).

Collectively, our results indicate that PYK2, but not FAK, regulates the cellular switch between fast and slow tumor cell motility within the primary tumor. Moreover, the lack of significant motility phenotype changes in cells knocked down for FAK suggests that PYK2 can compensate for FAK in regulating focal adhesion-mediated motility within primary xenograft tumors, but FAK cannot compensate for PYK2 in either focal adhesion-mediated motility or in invadopodia-dependent functions in these tumors.

### An integrated approach for elucidating the roles of PYK2 and FAK in breast cancer metastasis signaling

To elucidate the *in vivo* molecular and signaling mechanisms by which PYK2 and FAK regulate breast cancer invasiveness and metastasis, primary xenograft tumors were isolated from mice injected with control, PYK2-KD2, or FAK-KD1 breast cancer cells and subjected to RNA sequencing and mass spectrometry (**Figure 6A**). To elucidate the long-term changes following depletion of PYK2 or FAK in breast tumors, we performed enrichment analysis of differentially expressed genes (**Figure 6B and Supplementary Tables 1, 2**). The analysis revealed that the adhesion and P21 Activated Kinase 1 (PAK) pathways were up-regulated in both PYK2- and FAK-KD, while the Phospholipase C (PLC) pathway was down-regulated in both PYK2- and FAK-KD cells. The Integrin pathway was oppositely regulated in PYK2-KD (down) and FAK-KD (up) tumors. Importantly, the ECM remodeling pathway was down regulated in PYK2-KD tumors and up-regulated in FAK-KD tumors, further supporting our previous *in vitro* observations^43^ and our *in vivo* data presented herein. Interestingly, reduced actin nucleation by the ARP-WASp complex and stem cell differentiation were observed in FAK-KD tumors only.

**Figure 6.**
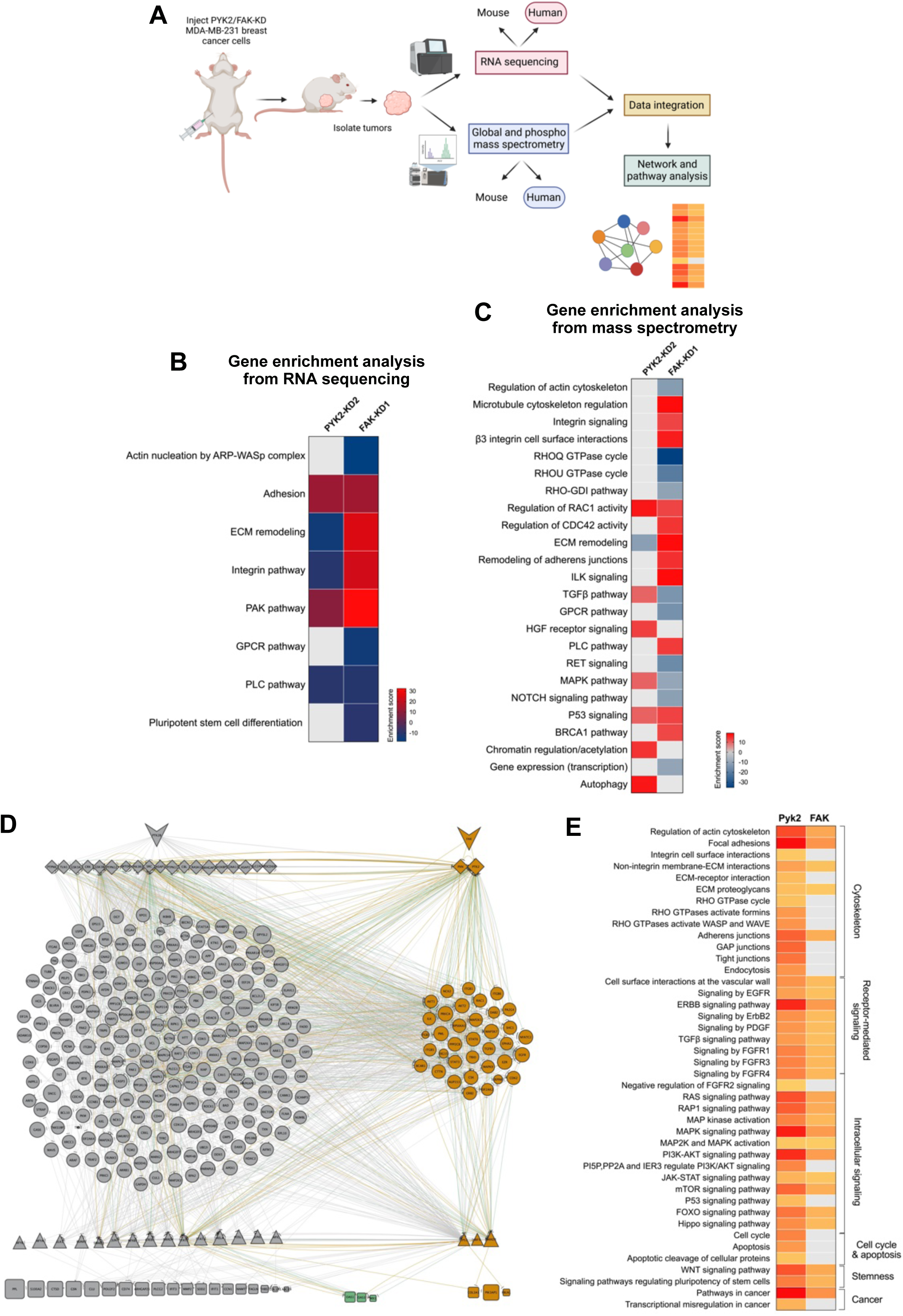
Integrated data analysis of PYK2-KD and FAK-KD xenograft mammary tumors. **(A)** A cartoon description of the transcriptomics/proteomics experiment from control, PYK2-KD, and FAK-KD mammary xenograft tumors. **(B)** Pathway enrichment analysis of transcriptomics in PYK2-KD2 and FAK-KD1 tumors. Differentially expressed genes in both tumors were used for the analysis. Gray, N/A. **(C)** Enrichment analysis of pathways in PYK2-KD and FAK-KD tumors. Differentially expressed total, phospho-tyrosine, and phospho-serine/threonine proteins were used for the analysis. Gray, N/A. **(D)** Merged network containing both the PYK2 network (left) and the FAK network (right). The shape of the nodes indicates their association with a specific layer (PYK2/FAK, PPI, or TF-DEG layer). Nodes and edges belonging to FAK, PYK2, or both causal networks are labeled in orange, grey, or green, respectively. The size of the nodes is proportional to the difference in protein abundance between PYK2 *vs.* FAK. **(E)** Enrichment analysis of the PYK2 and FAK causal networks; pathway values are expressed as experimental mean.

To gain an insight into the role of PYK2 and FAK in the immediate short-term signaling events that occur at the protein level in primary xenograft tumors, we performed enrichment analysis on the combined proteomics data from global, phosphotyrosine, and phosphoserine/threonine mass spectrometry (**Figure 6C and Supplementary Tables 1, 2**). Examination of the results revealed that both PYK2 and FAK have a significant increase in the regulation of RAC1-mediated pathway and in P53-mediated signaling pathway. Similar to what was observed in our RNA sequencing analysis above, and supporting our previous *in vitro* and current *in vivo* data, the ECM remodeling pathway is up regulated for FAK-KD and down regulated for PYK2-KD. The MAPK-mediated signaling pathway was regulated similarly to the ECM remodeling pathway. Interestingly, only FAK-KD tumors showed changes in actin cytoskeleton pathway (down), microtubule cytoskeleton regulation pathway (up), integrin-mediated signaling pathway (up), and in BRCA1-mediated signaling pathway (up). Only PYK2-KD tumors showed changes in autophagy signaling pathway (up) and in chromatin regulation/acetylation pathway (up).

To gain further insight into the signaling pathways and networks regulated by PYK2 and FAK, we integrated proteomics and transcriptomics data using *a priori* knowledge of human protein-protein interactions (PPIs) and regulatory interactions (transcription factor (TF) - target gene (TG)), taken from OmniPath and DoRothEA databases, respectively. Proteomics data were used to find expressed genes in the PYK2-KD and FAK-KD tumors. Transcriptomics data were also used to find differentially expressed genes in the knockdown *vs.* control tumors. Upstream interactions between FAK or PYK2 and their interacting human binding proteins were identified by filtering for condition-specific PPIs stemming from PYK2 and FAK. Condition specific regulators of differentially expressed genes (DEGs) were also identified by filtering the condition-specific TF-TG interactions for regulators targeting DEGs in PYK2/FAK-KD *vs.* control. Separate networks were built to construct the signal in PYK2-KD tumors, where FAK is present (FAK network), or in FAK-KD where PYK2 is present (PYK2 network). Comparison of the PYK2 network to the FAK network revealed 53 common proteins within the PPI layer, 186 proteins unique to Pyk2 only, and 27 proteins unique to FAK only (**Figure 6D, Supplementary Figures 2, 3, 4 and Supplementary Table 3**). In the regulatory network layer, three TFs appeared as unique to FAK (TP53, IRF9, SMAD3) and 15 TFs were unique to PYK2. Accordingly, 19 DEGs appeared in the PYK2 network and three (*COL3A1, PIK3AP1, FBLN1*) appeared in the FAK network. Three DEGs (*OAS1, OAS3, MX1*) appeared as common DEGs for both PYK2 and FAK.

Next, functional enrichment analysis of the PYK2 and FAK causal networks was performed using Adaptive NUll distriButIon of X-talk (ANUBIX)^51^ to investigate altered signaling pathways driven by FAK in PYK2-KD tumors or by PYK2 in FAK-KD tumors (**Figure 6E and Supplementary Table 3**). Among the enriched signaling pathways, some processes, such as regulation of actin cytoskeleton, EGFR-, PDGFR-, FGFRs-, and MAPK-mediated signaling, JAK-STAT-mediated signaling, PI3K-AKT-mediated signaling, mTOR-mediated signaling, WNT-mediated signaling, and pluripotency of stem cells were significantly changed in both PYK2 and FAK KD tumors. Interestingly, the focal adhesion pathway was also enriched in both PYK2 and FAK networks, further supporting our previous *in vitro* results showing that both PYK2 and FAK localize to focal adhesions and regulate tumor cell motility^43^ and supporting our intravital imaging results presented above, showing that PYK2 regulates focal adhesion-mediated fast motility and can compensate for FAK in regions far from blood vessels. Other signaling pathways such as RHO GTPase-mediated signaling, ECM-receptor interactions, endocytosis, and apoptosis were unique to PYK2 in this analysis. Together, our transcriptomics, proteomics, and integrated data analyses validate the *in vivo* results showing that both PYK2 and FAK are involved in tumor progression and in cancer cell motility and invasiveness, while uncovering new signaling pathways regulated by PYK2 and FAK in xenograft breast tumors.

### PYK2 and FAK coordinate the balance between focal adhesions and invadopodia in breast tumors

We have previously demonstrated that PYK2 and FAK modulate breast tumor cell invasiveness by coordinating the balance between focal adhesion-mediated migration and invadopodia-dependent invasion^43,44^. While FAK localizes only to focal adhesions (where it mediates Src-dependent focal adhesion dynamics and subsequent cell migration), PYK2 localizes to both invadopodia (where it mediates their maturation by activating the Src-Arg-cortactin axis) and to focal adhesions (where it regulates cell motility^43,44^). Coordination between both invadopodia-mediated invasion and focal adhesion-mediated motility is necessary for breast cancer cell intravasation and subsequent metastatic dissemination. To explore the *in vivo* role of PYK2 and FAK in focal adhesion mediated motility and invadopodia mediated invasion, we used lists of manually curated invadopodia and focal adhesion proteins (**Supplementary Table 4**) and built invadopodia and focal adhesion sub-networks from the PYK2 and FAK causal networks, to visualize how specific altered signaling pathways that are driven by FAK in PYK2-KD or by PYK2 in FAK-KD affect downstream invadopodia or focal adhesion-related functions.

Examination of the PYK2 and FAK networks revealed that both are enriched for invadopodia proteins with a higher significance for PYK2 network (PYK2 network: *p* = 2.7e^-25^; FAK network: *p* = 1.6e^-17^) (**Figure 7A, B and Supplementary Table 4**). Additionally, the total number of invadopodia nodes was higher for PYK2 (80 nodes) than for FAK (35 nodes). Overlap of the proteins in both groups revealed that all FAK proteins are included in the PYK2 invadopodia list, but PYK2 has additional 45 invadopodia proteins which do not appear in the FAK network (**Figure 7C**). These data support the hypothesis that PYK2 has a critical role in invadopodia regulation. Both PYK2 and FAK networks were also enriched for focal adhesion proteins (PYK2 network: *p* = 1.8e^-5^; FAK network: *p* = 1.5e^-9^) (**Figure 7D, E and Supplementary Table 4**). Importantly, the total number of focal adhesion nodes was higher in PYK2 (172 nodes) than in FAK (68 nodes) and overlap of both groups revealed they have 68 common focal adhesion nodes, but PYK2 has additional unique 104 nodes that do not appear in the FAK network (**Figure 7F**). These results suggest that PYK2 plays a role similar to FAK in focal adhesions, but also has additional roles. All in all, these data support our hypothesis (which is based on intravital imaging results) that PYK2 can compensate for FAK in focal adhesions, but FAK cannot compensate for PYK2 in either invadopodia or in focal adhesions.

**Figure 7.**
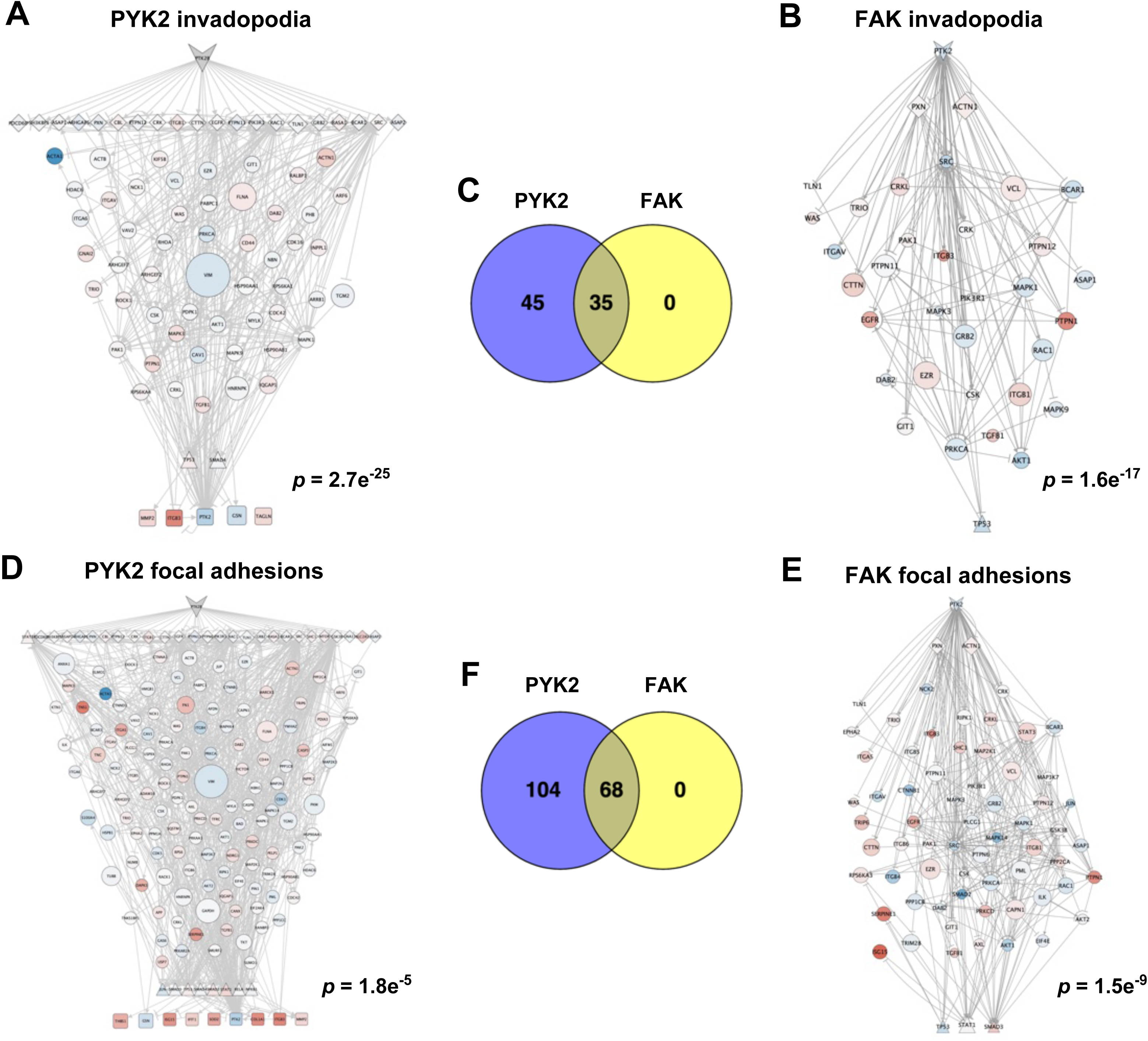
Network analysis of PYK2 and FAK in invadopodia and focal adhesion regulation. **(A)** Invadopodia sub-network of PYK2. *p* = 2.7e^-25^, demonstrating that the signaling network of PYK2 is enriched in invadopodia proteins. **(B)** Invadopodia sub-network of FAK. Invadopodia enrichment score: *p* = 1.6e^-17^. **(C)** Overlap of PYK2 and FAK invadopodia signaling proteins. PYK2, 80 proteins; FAK, 35 proteins; overlap: 35 proteins. **(D)** Focal adhesion sub-network of PYK2. Enrichment score: *p* = 1.8e^-5^. **(E)** Focal adhesion sub-network of FAK. Enrichment score: *p* = 1.5e^-9^. **(F)** Overlap of PYK2 and FAK focal adhesion signaling proteins. PYK2, 172 proteins; FAK, 68 proteins; overlap, 68 proteins.

## DISCUSSION

Accumulating evidence suggests that both PYK2 and FAK play a role in breast cancer progression and metastasis, but the *in vivo* cellular and signaling mechanisms by which they do so are poorly understood at present. Here, we show that knockdown of FAK, and to a lesser extent PYK2, reduces primary tumor size due to reduced proliferation in FAK-KD tumors and increased apoptosis in PYK2-KD tumors. Despite the observation that PYK2 and FAK oppositely regulate invadopodia-like protrusions and ECM degradation within the primary tumors, ablation of either kinase significantly reduced the number of circulating tumor cells and disseminated tumor cells in tumor bearing mice. This phenotype can be explained by reduced TMEM-doorway activity which leads to decreased intravasation in both PYK2-KD and FAK-KD tumors. Using single cell resolution intravital imaging of primary knockdown tumors, we describe, for the first time, a role for PYK2 in regulating a motility phenotype switch between focal adhesion-mediated fast motility in regions far from blood vessels and invadopodia-dependent ECM degradation-mediated slow motility in regions close to tumor blood vessels (**Table 1**). Additionally, integrated transcriptomics, proteomics, and protein-protein interaction data analysis support these intravital imaging results, while exposing new and novel signaling pathways by which PYK2 and FAK regulate tumorigenesis and metastasis (**Supplementary Table 5**). Our work identifies PYK2 and FAK as novel mediators of tumor progression and metastasis and elucidates, for the first time, the *in vivo* cellular and signaling mechanisms by which they do so. Moreover, although highly homologous, knockdown of either PYK2 or FAK significantly reduced breast cancer invasiveness and intravasation, indicating that these two kinases regulate progression and metastasis *via* distinct mechanisms.

**Table 1.** Summary of the *in vivo* phenotypes of PYK2-KD and FAK-KD xenograft tumors.

| ***In vivo* phenotype** | **PYK2-KD** | **FAK-KD** | **Figure** |
| --- | --- | --- | --- |
| Primary tumor size | Decreased | Decreased | 2 |
| *In vivo* invasion (needle collection assay) | Decreased | Decreased | 3 |
| CTCs | Decreased | Decreased | 3 |
| Disseminated tumor cells from orthotopic primary tumors | Decreased | Decreased | 3 |
| TMEM doorway assembly | No change | No change | 4 |
| TMEM doorway activation | Decreased | Decreased | 4 |
| *MENA*<sup>INV</sup> expression | Decreased | No change | 4 |
| Localization to invadopodia-like structures | Yes | No | 5 |
| Tumor cell protrusions | Decreased | Increased | 5 |
| *In vivo* MMP activity | Decreased | Increased | 5 |
| Tumor cell speed close/far from blood vessels | Increased/decreased | No change | 5 |

Using intravital imaging of mammary tumors in live mice, two motility phenotypes of tumor cells, differing in speed of locomotion, were previously detected within the primary tumor: focal adhesion-mediated fast motility on ECM fibers, and slow invadopodia-mediated ECM degradation-associated motility. These distinct motility phenotypes are context-dependent and determined by a combination of cues present in the local microenvironment, among them collagen fiber density, tumor cell and macrophage density, and proximity of tumor cells to blood vessels, where they receive chemical and mechanical cues that induce invadopodia formation and activation. These two motility phenotypes both contribute to metastasis; tumor cells use fast locomotion to translocate to perivascular regions where they stop, due to ECM composition and stiffness, and switch to an invadopodium-associated phenotype that results in their intravasation and dissemination^34,42^. Along these lines, a recent study by Lengrand *et al.* suggests that PYK2 regulates the dynamic transition between focal adhesion-mediated migration and invadopodia-dependent invasion in invasive melanoma cell lines^52^. They also suggest that, because focal adhesions and invadopodia share many structural components, the dynamic exchange between these two structures, which is controlled by PYK2, regulates the cycles of migration and ECM-mediated invasion which are necessary for efficient cancer metastasis. Our intravital imaging experiments of breast cancer cell behavior within primary tumors support these results, and indicate that PYK2, but not FAK, regulates the motility phenotype switch between focal adhesion-mediated fast motility in regions far from blood vessels and invadopodia-mediated ECM invasion in regions close to blood vessels, where TMEM-doorway assembly and activation and subsequent intravasation take place.

We have previously shown that PYK2 localizes to invadopodia but can also co-localize with FAK to focal adhesions in breast cancer cells^43^. Moreover, depletion of either PYK2 or FAK in breast cancer cells significantly decreases their motility in 2D *in vitro* cultures, suggesting that the two kinases cannot compensate for each other and that, at least *in vitro*, they have different roles in focal adhesion-mediated motility. However, our intravital imaging results suggest that PYK2 can compensate for FAK in focal adhesion-mediated fast motility, but FAK cannot compensate for PYK2 in either focal adhesion-mediated motility or in invadopodia-mediated ECM invasion. These findings were further confirmed by our integrated data analysis which indicates that FAK interacting proteins at focal adhesions overlap with PYK2 interacting proteins completely, while PYK2 has additional interacting proteins. These results suggest that *in vivo*, PYK2 can compensate for FAK in focal adhesions but also has additional unique roles in these structures. Moreover, the fact that PYK2 can compensate for FAK in focal adhesion-mediated fast motility in regions far from blood vessels within primary tumors suggests that this function is indispensable for tumor cell invasiveness and for metastatic dissemination. The significant differences that exist between the focal adhesion-mediated motility behavior of PYK2-KD and FAK-KD cells *in vitro* versus *in vivo* may result from the fact that motility phenotypes of cancer cells within the primary tumor are determined by a combination of microenvironmental conditions, which do not exist in tissue culture.

The transcriptomics, proteomics, and integrated data presented in this paper support previous findings from our group and others and support the *in vivo* phenotypes described in this paper. They also reveal previously unexplored pathways by which PYK2 and FAK regulate tumorigenesis and metastasis, including apoptosis, autophagy, chromatin regulation, and DNA repair. While the smaller size of FAK-KD tumors may result from decreased tumor cell proliferation, our immunohistochemistry, transcriptomics, and integrated data showed increased apoptosis in PYK2-KD tumors, which involves changes in apoptosis-related genes and proteins such as Caspase 3, BAD, BAX, XIAP, FADD, and MAPK8 which may explain their smaller size. While Burdick *et al*. previously reported that PYK2 is involved in Benzo(a)pyrene (BaP) induced anti-apoptotic signaling (which promotes its carcinogenic effect on cultured mammary cells^53^), a role for PYK2 in inhibiting the apoptosis of cancer cells within the primary tumor has not been described to date. The specific role and signaling pathway which mediates inhibition of apoptosis by PYK2 needs further research and is a subject for future investigation.

Our *in vivo* results support previous *in vitro* data^43,52^ and show that knockdown of PYK2 decreases, while knockdown of FAK increases, invadopodia-like protrusion formation in tumor cells and subsequent tumor ECM degradation. Despite these higher activities both *in vitro* and in tumors, FAK-KD cells showed less invasiveness *in vitro*^43^ and reduced intravasation and lung metastasis *in vivo*. While the lower *in vitro* invasion of FAK-KD tumor cells could be explained by reduced focal adhesion-mediated motility on two-dimensional substrates, we did not observe any changes in the motility of these cells within tumors, either in regions far from blood vessels, where motility is mediated by focal adhesions, or in regions close to blood vessels, where motility is mainly mediated by invadopodia. Lower blood burden and dissemination of FAK-KD tumors could therefore be explained by decreased TMEM-doorway activity, which may result from reduced responsiveness of tumor cells to NOTCH1-mediated signaling in TMEM-doorways or by a decrease in other interactions with TMEM-doorway associated cells. The involvement of FAK in NOTCH1-mediated signaling of breast cancer is largely unknown, but some evidence exists in gastric cancer. Specifically, down-regulation of NOTCH1 inhibits human gastric cancer cell invasiveness through the activation of PTEN and de-phosphorylation of FAK and AKT^54^. Moreover, NOTCH1 mediated induction of miR151-3p and miR151-5p in these cells promotes viability, migration, invasion, and consequent progression of gastric cancer by increasing FAK expression^55^. Whether FAK is involved in NOTCH1 signaling in breast cancer in general and in TMEM doorway activation in particular by using the same mechanisms as in gastric cancer or by stabilization of N1ICD, similarly to PYK2, is a subject for future investigation.

The involvement of PYK2 in NOTCH1 signaling was recently described by Muller *et al*., who suggested that PYK2 stabilizes N1ICD (the intracellular cleaved form of NOTCH1) in the cytoplasm of breast cancer cells. Stabilized N1ICD then enters the nucleus, where it induces the transcription of *CCL2*, which is used by the tumor cells to attract macrophages into the tumor^56^. Indeed, *MENA^INV^* has been found to be one of the genes which is expressed in breast tumor cells following NOTCH1 signaling activation by TMEM doorway-associated macrophages^34,50,57^, and our data showed reduced expression of *MENA^INV^* in PYK2-KD tumors, suggesting that decreased TMEM doorway activation in these tumors may result from ablation of NOTCH1-mediated signaling to result in decreased *MENA^INV^*expression. *MENA^INV^* expression has been linked to TMEM doorway-associated trans-endothelial migration resulting in tumor cell intravasation, and extravasation at distant sites, making it an essential player in tumor cell metastatic dissemination^34,38,39,49^.

Despite their opposite activities, knockdown of either PYK2 or FAK significantly inhibits breast cancer dissemination in immune-compromised mice due to inhibition of TMEM doorway activation and subsequent tumor cell intravasation into the bloodstream. This result further implies that inhibition of PYK2 and/or FAK could block breast cancer metastasis. Considering the fact that PYK2, but not FAK, localizes to both focal adhesions and invadopodia and regulates the switch between the two motility phenotypes that control metastasis, it may be more beneficial to specifically inhibit PYK2 in tumors. Moreover, since FAK is ubiquitously expressed and the expression of PYK2 is limited, specific inhibition of PYK2 may cause fewer side effects and may therefore be more beneficial in blocking breast cancer metastasis. Unfortunately, no specific inhibitor for PYK2 exists at present and while dual PYK2/FAK inhibitors have been evaluated in combination therapy clinical trials of various cancers, these have only focused on reducing primary tumor size^58,59^, and have not evaluated the impact on dissemination. Once developed, a specific PYK2 inhibitor will be ideally situated to block the metastatic spread of breast cancer cells when used in combination with standard cytotoxic therapies that prevent the growth of tumor cells within the primary tumor and in secondary and tertiary metastatic sites.

## ACKNOWLEDGEMENTS

We wish to thank Dr. Meital Kupervaser and Dr. Yishai Levin from G-INCPM at Weizmann Institute for technical assistance and protocol development of tumor mass spectrometry. We are grateful to the Gruss Lipper Biophotonics Center and Analytical Imaging Facility at Albert Einstein college of Medicine for their assistance in imaging and to the Einstein Histotechnology and Comparative Pathology Facility for their excellent histology service. This work was funded by the Israel Cancer Research Fund (grant number 20-101-PG), the Israel Cancer Association (grant number 20210071) and the Israel Science Foundation (grant number 2142/21) (to H. Gil-Henn), and CA255153, CA257885, the Spatz Family Foundation, and Evelyn Gruss Lipper Charitable Foundation (to G. S. Karagiannis, D. Entenberg, M. H. Oktay and J. S. Condeelis). The work of T. Korcsmaros and M. Poletti was supported by the UKRI BBSRC Gut Microbes and Health Institute Strategic Program BB/R012490/1 and its constituent projects BBS/E/F/000PR10353 and BBS/E/F/000PR10355, as well as a UKRI BBSRC Core Strategic Program Grant for Genomes to Food Security (grant number BB/CSP1720/1) and its constituent work packages, BBS/E/T/000PR9819 and BBS/E/T/000PR9817. MP was also supported by the BBSRC Norwich Research Park Biosciences Doctoral Training Partnership (grant number BB/M011216/1).

## MATERIALS AND METHODS

### Proteogenomic analysis

Data from two cohorts of 1,080 and 825 breast invasive carcinoma samples was obtained from The Cancer Genome Atlas (TCGA, https://cancergenome.nih.gov)^60^ and analyzed with cBioPortal tools (http://www.cbioportal.org)^61^ using MATLAB. The analyzed datasets contain mRNA-seq expression Z-scores (RNA-Seq V2 RSEM) and Agilent microarray mRNA Z-scores computed as the relative expression of an individual gene and tumor to the expression distribution of all samples that are diploid for the gene. Copy number alterations were collected from both cohorts and calculated using genomic identification of significant targets in cancer (GISTIC), a statistical method that derives a scoring metric incorporating both the magnitude and frequency of copy number changes at every genomic position^62^. GISTIC is an algorithm that attempts to identify significantly altered regions of amplification or deletion and uses discrete copy number calls: homozygous deletion; heterozygous loss; neutral; gain; high amplification. Breast cancer samples are annotated with OS and/or DFS time and censorship status. Samples were split into high and low expressing groups based upon median mRNA expression Z-scores. Associations between Z-scores and patient survival (DFS and OS) were assessed by Kaplan–Meier time-event curves and Mantel-Haenszel hazard ratios using an implementation of Kaplan–Meier log rank testing from MATLAB Exchange. All statistical tests were two-sided. Mass-spectrometry based proteomic characterization of 102 breast cancer tumor samples^63^ was obtained from the Clinical Proteomic Tumor Analysis Consortium (CPTAC) Data Portal (https://cptac-data-portal.georgetown.edu)^64^ with cBioPortal using MATLAB. For each protein target, Z-scores were determined, and associated P-values were calculated across all samples.

### Distant metastasis free survival analysis

Microarray datasets with Distant Metastasis Free Survival (DMFS) annotation were obtained from the NCBI Gene Expression Omnibus (GEO) data repository^65^ of high-throughput microarray experimental data. The meta-cohort dataset comprised of 1,650 tumor expression profiles of primary invasive breast cancer based on the Affymetrix U133 GeneChip microarray platform. Queried transcripts included 22,283 probe sets common to all microarrays in all study populations. Assembly of the datasets was performed using MATLAB (The Mathworks, Inc.) and GEO series (GSE) files were extracted via GEO accessions GSE11121, GSE25055, GSE7390, GSE25065, GSE17705, GSE12093, GSE1456, GSE5327 and GSE45255. The datasets have been retrieved with uniform normalization of probe intensities with MAS 5.0^66^ using global scaling with a trimmed mean target intensity of each array arbitrary set to 600. Cross-population batch effects were corrected using Z-score transformation ^67^.

The tumor profiles represent primary invasive breast tumors sampled at the time of surgical resection, annotated with DMFS time and censorship status. Patient samples were split into high and low expressing groups based upon median gene expression. Associations between normalized gene expression and patient survival (DMFS) were assessed by Kaplan-Meier time-event curves and Mantel-Haenszel hazard ratios using an implementation of Kaplan-Meier log rank testing from MATLAB Exchange. All statistical tests were two-sided. The synergy index (SI) was calculated as a means of evaluating additive interaction^68,69^. The synergy index can be interpreted as the excess risk from overexpression of both genes relative to the risk of overexpression of the genes separately. SI of 1 indicates no synergism, and an SI >1 indicates synergistic interaction between the two genes.

### Constructs and cell lines

MDA-MB-231 human breast adenocarcinoma cells and HEK293T cells were from the American Type Culture Collection. GP2 packaging cell line was from Clontech. MDA-MB-231 cells stably expressing Dendra 2 (MDA-MB-231/Dendra2 cell line) were previously described^42^. For shRNA-mediated knockdown in MDA-MB-231/Dendra2 cells, the following sequences were cloned into the pLKO.2 retroviral plasmid: PYK2-KD1 (5’-GGTGGTGGTACCAGTAGAT-3’), PYK2-KD2 (5’-GGATCATCATGGAATTGTA-3’), FAK-KD1 (5’-GGGCATCATTCAGAAGATA-3’), FAK-KD2 (5’-TAGTACAGCTCTTGCATAT-3’). Stable knockdown cell lines were generated by transfecting pLKO.2 viral expression vectors carrying shRNA sequences along with pCMV-dR8.2 and pCMV-VSV-G plasmids into HEK293T cells. Viral sups were used to infect MDA-MB-231/Dendra2 cells, which were then selected with 10 μg/ml puromycin. A control cell line was also generated by transfection of empty vector using the same protocol. Knockdown or overexpression levels were verified by immunoblotting. MDA-MB-231 cells stably expressing both cortactin-TagRFP and either PYK2-GFP or FAK-GFP were generated by infecting the cortactin-TagRFP cell line^70^ with viral sups that were collected from GP2 cells transfected with pQCTK-PYK2-GFP, or pQCTK-FAK-GFP, respectively, followed by selection with 200 μg/ml hygromycin B.

### Immunoblotting

Immunoblotting analysis was performed as previously described^40^. Briefly, cells were washed in cold phosphate-buffered saline (PBS) and lysed in modified RIPA buffer (50mM Tris pH 7.2, 150mM NaCl, 1% NP-40, 0.5% deoxycholate, 0.1% SDS, 1mM ethylenediamine tetra acetic acid, 2mM NaF, 1mM Na3VO4 and protease inhibitors). Samples were resolved by SDS-PAGE, transferred to nitrocellulose, blocked in Odyssey blocking solution (LI-COR Biosciences), incubated with primary antibodies (anti-PYK2 #3292; Cell Signaling Technology, anti-FAK 610088; BD Biosciences, anti-β-actin A5441; Sigma-Aldrich) overnight at 4°C and then with secondary antibodies (LiCOR) for one hour at room temperature. Blots were imaged using the Odyssey CLx imaging system.

### Mouse xenograft model

All experimental procedures were conducted in accordance with the Federation of Laboratory Animal Science Associations (FELSA) and the National Institutes of Health regulations and were approved by the Bar-Ilan University and the Albert Einstein College of Medicine animal care and use (IACUC) committees. Mouse xenograft tumors containing control, PYK2-KD, or FAK-KD cell lines were generated by injecting a total of 2×10^6^ MDA-MB-231/Dendra2 cells resuspended in 20% collagen I (BD Biosciences) in PBS into the lower left mammary gland of SCID mice. For tumor growth experiments, samples were harvested 10 weeks following injection. All other DTC, intravital imaging, transcriptomics, and proteomics experiments were performed on tumors that were 1-1.2 cm in diameter. In experiments described in Figures 3, 4B, 4D, 5, 6, 7, 9, 10, and Supplementary Figures 1, 2, 3 tumors generated from PYK2-KD2 and FAK-KD1 were used.

### Tumor immunohistochemistry

Tumors were excised at 1-1.2 cm diameter, fixed in formalin and paraffin embedded. Sections from the middle of the primary tumors were stained with hematoxylin and eosin for general histology, or immunostained with specific antibodies: anti-Ki67 (VP-K451, Vector Laboratories) for proliferation or anti-cleaved caspase 3 (9661, Cell Signaling Technology) for apoptosis. Briefly, samples for immunohistochemistry were sectioned at 5 mm and deparaffinized in xylene followed by graded alcohols. Antigen retrieval was performed in 10mM sodium citrate buffer at pH 6.0, heated to 96°C, for 20 min. Endogenous peroxidase activity was quenched using 3% hydrogen peroxide in PBS for 10 min. Blocking was performed by incubating sections in 5% normal donkey serum with 2% bovine serum albumin for 1 h. Tumor sections were stained by routine immunohistochemistry methods, using HRP rabbit polymer conjugate (Invitrogen), for 20 min to localize the antibody bound to antigen, with diaminobenzidine as the final chromogen. All immunostained sections were lightly counterstained with hematoxylin. Proliferation and apoptosis were quantified by counting the Ki67 positive cells or cleaved caspase 3 positive cells (brown) over total number of cells (blue) in five representative fields per tumor (at 40X) and a total of three tumors per group. Necrotic tumor areas were excluded from the analysis.

### *In vivo* invasion assay

33G needles were filled with Matrigel and L15-BSA with or without (control) addition of 25 nM human recombinant EGF (Invitrogen). Mice were anesthetized using 5% isoflurane and laid on their back. The isoflurane was reduced to 2%, and a small patch of the skin over the tumor was removed. Six 25G needles with blocking wires were inserted to a depth of 2 mm from both sides of the tumor. Blocking wires were then removed, and one Matrigel-filled 33G needles (either with or without EGF) was inserted into each guide needle. The needles were left in the tumor for four hours. Isoflurane concentration was slowly lowered to 0.5% during experiment to keep the mouse breathing even and unlabored. Following four hours of collection, the needles are removed, and the total number of cells collected was determined by 40,6-diamidino-2-phenylindole (DAPI) staining.

### Quantification of intravasated circulating tumor cells

Mice were anesthetized with isoflurane and blood was drawn from the right ventricle by heart puncture using heparin-coated 25G needles. Erythrocytes were lysed using 10 ml of 1xRBC lysis buffer (eBioscience). Samples were centrifuged at 1,000 rpm for 5 min, then cell pellets were reconstituted in 10 ml DMEM/F12 supplemented with 20% FBS and plated in 10 cm petri dishes. Following cell attachment to the dish, single tumor cells were counted. The total number of tumor cells counted was normalized to the volume of blood collected for each mouse as shown in **Figure 4D**.

### Spontaneous lung metastasis assay

Disseminated single tumor cells and micro-metastases were measured in SCID mice bearing orthotopic control, PYK2-KD or FAK-KD MDA-MB-231/Dendra2-derived tumors of equal size (1-1.2 cm in diameter). Five minutes before sacrifice, mice were injected intravenously with rhodamine-conjugated lectin (Vector Laboratories) for visualizing blood vessels. The largest lung lobe of each mouse was imaged using an Olympus FV1000-MPE multiphoton microscope (Olympus, Center Valley, PA, USA) and single extravascular Dendra2 fluorescent cancer cells or micro metastases were counted.

### TMEM doorway immunohistochemistry

Mice were sacrificed and mammary tumors were extracted and immersed in 10% formalin in a volume ratio of tumor to formalin 1:7. Tumors were fixed for 24-48 hours, embedded in paraffin, and processed for histological examination. Tumors were sliced in sections of 5 μm and stained for TMEM doorways with anti-endomucin (SC-65495; Santa Cruz Biotechnology), Iba1 (019-19741; Wako) and anti-pan Mena (510693; BD Biosciences). Areas containing invasive tumor tissue suitable for TMEM doorway analysis were identified by low-power scanning using the following criteria: high density of tumor cells, adequacy of tumor, lack of necrosis or inflammation, and lack of artefacts such as retraction or folds. The assessment of TMEM doorway scores was performed with Adobe Photoshop on five high power (440x330 µm) digital images of the most representative areas of the tumor. TMEM doorways were scored manually by a pathologist in a blinded fashion. The total count of TMEM doorways for each image was tabulated, and the scores from all five images were summed to give the final TMEM doorway score for each mouse tumor, expressed as the number of TMEM doorways per total area.

### Vascular opening

One hour before sacrifice, 3 mg of 155 kDa TMR-dextran was administered by tail vein injection to label sites of vascular opening. Assessment of vascular opening was performed using multi-immunofluorescence in an FFPE section with a sequential TMEM doorway triple-IHC section which was already labeled as above, as previously described^48^. Following standard slide preparation, slides were immunolabeled with anti-endomucin (sc-65495, Santa Cruz Biotechnology) and anti-TMR (A6397; Life Technologies). Slides were washed three times in 0.05% PBST and incubated with fluorescently labeled secondary antibodies (Alexa 488 donkey anti rabbit and Alexa 568 goat anti rat) for 60 minutes at room temperature. Slides were then washed three times in 0.05% PBST, incubated with DAPI for five minutes, and mounted using ProLong Gold antifade reagent (Life Technologies). The slides were imaged on Panoramic 250 Flash II digital whole slide scanner using a 20X 0.75 NA objective lens. In each image, the endomucin and TMR channels were each thresholded above background based on their intensity by using contrast adjustment. For each vascular profile, the endomucin channel was then used as an exclusion mask to the dextran channel, to directly designate an ROI that belonged exclusively to extravascular dextran. The sequential TMEM doorway IHC sections were then used to assess whether these leaky profiles had an associated TMEM doorway. To directly compare vascular opening between groups, the extravascular dextran ROI was expressed as an area fraction, and an average among all ROIs for each mouse was reported.

### Quantitative real-time PCR (qRT-PCR)

RNA was extracted from control and knockdown tumors using the RNeasy Mini kit (cat#74104, Qiagen). RNA was quantified using NanoDrop TM 2000 (Thermo Scientific). Two micrograms of total RNA were reverse transcribed using High-Capacity RNA to cDNA kit (cat#4387406, Applied Biosystems). Quantitative PCR analysis was performed using TaqMan Fast advanced master mix (cat#AB-4444557, Applied Biosystems) or FAST SYBR Green master mix (cat#AB-4385612, Applied Biosystems) in triplicates using the StepOnePlus real-time PCR system (Applied Biosystems) and 10 ng cDNA template. Results were quantified using the comparative Ct (ΔΔCt) method. Primer sequences used in this study are described in the table below.

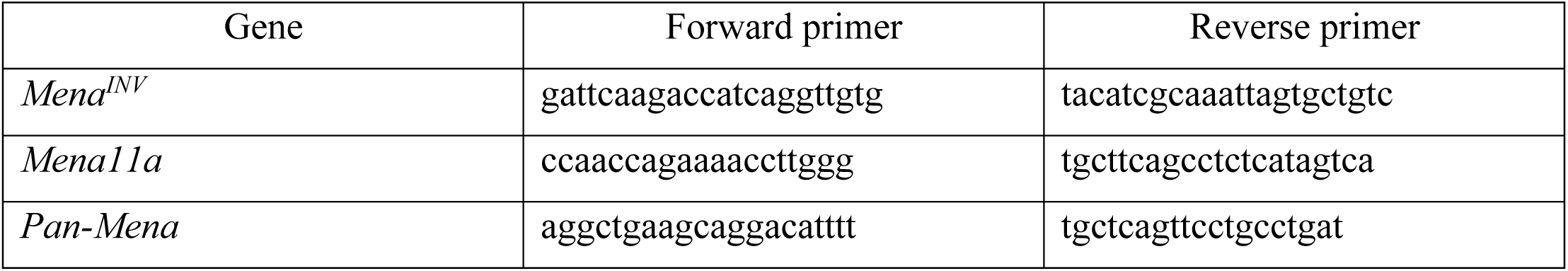

### NanoString analysis

NanoString analysis was performed on tumors collected at the same time (10 weeks following injection, 5 tumors per group) according to the manufacturer’s protocol. In brief, 150 ng total RNA from each sample were hybridized with reporter and capture probe sets (nCounter Tumor Signaling 360 panel) by incubating for 18 hours at 65°C. Following hybridization, samples were immobilized and then counted on the nCounter FLEX system. Data were analyzed using the nSolver 4.0 system and nSolver advanced data analysis module (2.0.143).

### Tumor immunostaining

Eight-week-old female SCID mice were injected with MDA-MB-231 cells. When tumors reached the size of 1-1.2 cm in diameter mice were sacrificed and tumors or lungs were excised, fixed in 4% paraformaldehyde overnight, washed for 1 hour in cold PBS and dehydrated overnight in 30% sucrose. Next, tissue was embedded in OCT and 5 μm thick cryostat sections were placed on silane-coated slides and dried at room temperature followed by permeabilization with 0.1% Triton X-100 for 10 minutes and blocking in 1% BSA and 1% FBS for 1 hour at room temperature. Samples were then incubated overnight at 4°C with the indicated primary antibodies, washed, and incubated with the appropriate secondary antibodies. Nuclei were counterstained with DAPI. Tissue was imaged using an inverted laser scanning confocal microscope (Zeiss LSM780; 60x, NA 1.4, oil objective, ZEN black edition acquisition software).

### *In vivo* MMP activity assay

Eight-week-old female SCID mice were injected with control, PYK2-KD, or FAK-KD MDA-MB-231/Dendra2 cells. When tumors reached the size of 1-1.2 cm in diameter, mice were tail vein-injected with of MMP Sense 645 FAST fluorescent imaging agent (4 nmol in 100 μl of PBS; NEV10100, Perkin-Elmer). Twenty-four hours following injection, mice were sacrificed, and tumors were excised and processed for immunofluorescence. Cleavage of MMP Sense 645 FAST was analyzed by quantifying the average fluorescent signal area in pixels per field using ImageJ.

### Multiphoton intravital imaging

SCID mice bearing tumors generated by MDA-MB-231/Dendra2 cells at size of 1.3 cm in diameter were injected with tetramethylrhodamine-labeled 155kD dextran (Sigma-Aldrich) ten minutes before imaging. A mammary imaging window was surgically implanted on the primary tumor. One day after the surgery, mice were placed on an inverted microscope and imaged continuously using a custom-built two-laser multi-photon microscope^71^ with 880 nm (for Dendra2 and dextran visualization) excitation for up to 6 hours. Random fields of 512×512 μm at 0.67-1 µm per pixel were imaged for a depth of 100 μm (21 slices at 5 μm steps) for a total of 30-60 minutes each. Images were analyzed in Fiji/ImageJ^72^.

### RNA sequencing and gene mapping

RNA from dissected tumors was purified using the RNeasy mini kit (Quigen) according to the manufacturer’s instructions. Integrity of isolated RNA was tested using the Agilent RNA Pico 6000 kit and Bioanalyzer 2100 at the Genome Technology Center of the Azrieli Faculty of Medicine, Bar-Ilan University. 350 ng of total RNA were used for mRNA enrichment using NEBNext mRNA polyA isolation module (NEB, #E7490L) and libraries for Illumina sequencing were prepared using the NEBNext Ultra II RNA kit (NEB, #E7770L). Quantification of the library was performed using dsDNA HS assay kit and Qubit (Molecular Probes, Life Technologies) and quantification was performed using the Agilent DNA HS kit and Bioanalyzer 2100. 10 nM of each library were pooled together and were diluted to 4 nM according to NextSeq manufacturer’s instructions. 1.1 pM was loaded into the flow cell with 1% PhiX library control. 75SR reads were sequenced on 8 lanes of an Illumina NextSeq 550.

Gene mapping was performed by The Weizmann Institute INCPM Bioinformatics unit. Briefly, poly A/T stretches and Illumina adapters were trimmed from the reads using cutadapt^73^, and resulting reads shorter than 30 bp were discarded. Reads were mapped to the H. *Sapiens* reference genome GRCh38 using STAR^74^, supplied with genome annotations downloaded from Ensembl (with EndToEnd option and outFilterMismatchNoverLmax set to 0.04). Expression levels for each gene were quantified using htseq-count^75^, using the gtf as above. Differentially expressed genes were identified using DESeq2^76^ with the betaPrior, cooksCutoff and independent filtering parameter set to false. Raw *p* values were adjusted for multiple testing using the procedure of Benjamini and Hochberg. Pipelines was run using snakemake^77^.

### Mass spectrometry

Dissected tumors were washed in PBS and homogenized at room temperature in urea lysis buffer (8 M urea, 40 mM Tris pH 7.6, 5% SDS supplemented with phosphatase inhibitor cocktails 2, 3 (Sigma P5726, P0044)). Lysates were centrifuged at 20,000 g for 60 minutes; supernatants were transferred into new tubes and snap frozen in liquid nitrogen. For global mass spectrometry analysis, samples were subjected to in-solution tryptic digestion followed by desalting. The resulting peptides were analyzed using nanoflow liquid chromatography (nanoAcquity) coupled to high resolution, high mass accuracy mass spectrometry (Q Executive HFX). Each sample was analyzed on the instrument separately in a random order in discovery mode. Raw data was processed using MaxQuant v1.6.0.16. The data was searched with the Andromeda search engine against the human and murine proteome databases appended with common lab protein contaminants and thee following modifications: carbamidomethylation of C as a fixed modification, oxidation of M, and protein N-terminal acetylation as variable ones. The LFQ (label-free quantification) intensities were calculated and used for further calculations using Perseus v1.6.0.7. Decoy hits were filtered out as well as proteins that were identified based on modified peptide only. The LFQ intensities were log transformed and only proteins that had at least 2 valid values in at least one experimental group were kept. The remaining missing values were imputed. A total of 5270 proteins were identified and quantified using this method.

For phospho-Ser/Thr mass spectrometry analysis, samples were subjected to enrichment of phosphorylated proteins using the Agilent automated robot with IMAC tips following in-solution tryptic digestion and desalting. The resulPng pepPdes were analyzed using nanoflow liquid chromatography (nanoAcquity) coupled to high resoluPon, high mass accuracy mass spectrometry (Q ExacPve HFX). Each sample was analyzed on the instrument separately in a random order in discovery mode. Raw data was processed with MaxQuant v1.6.6.0. The data was searched with the Andromeda search engine against the human and murine proteome databases appended with common lab protein contaminants and the following modificaPons: CarbamidomethylaPon of C as a fixed modificaPon, oxidaPon of M, phosphorylaPon of STY and protein N-terminal acetylaPon as variable ones. The phospho-sites intensiPes were calculated and used for further calculaPons using Perseus v1.6.2.3. Decoy hits were filtered out. The sites intensiPes were log transformed and only sites that had at least 3 valid values in at least one experimental group were kept. The log-transformed intensiPes were normalized by subtracPng the median. A total of approximately 13,000 phospho-sites were identified using this method.

For phospho-Tyr mass spectrometry analysis, samples were subjected to enrichment of tyrosine phosphorylated proteins using an the PTMScan HS phopshotyrosine kit (Cell Signaling Technology, #38572) following in-solution tryptic digestion and desalting. The resulLng pepLdes were analyzed using nanoflow liquid chromatography (nanoAcquity) coupled to high resoluLon, high mass accuracy mass spectrometry (Q ExacLve HFX). Each sample was analyzed on the instrument separately in a random order in discovery mode. Raw data was processed with MaxQuant v1.6.6.0. The data was searched with the Andromeda search engine against the human and murine proteome databases appended with common lab protein contaminants and the following modificaLons: CarbamidomethylaLon of C as a fixed modificaLon, oxidaLon of M, phosphorylaLon of STY and protein N-terminal acetylaLon as variable ones. The phosphosites intensiLes were calculated and used for further calculaLons using Perseus v1.6.2.3. Decoy hits were filtered out. The sites intensiLes were log transformed and only sites that had at least 3 valid values in at least one experimental group were kept. The log-transformed intensiLes were normalized by subtracLng the median. A total of 415 pTyr sites were identified using this method.

### Enrichment analysis

Differentially expressed genes were filtered for adjusted *p*-value (padj) ≤ 0.05, Log2FC > Ι 1Ι, and max count > 30. Differentially expressed proteins were defined according to the following conditions: 0.5 > ratio > 2, difference < −1 or > 1, *p* ≤ 0.05, 2 or more unique peptides. Peptides which are derived from human proteins or mouse/human proteins were included. Transcriptomics and proteomics enrichment analyses were performed using GeneAnalytics (https://geneanalytics.genecards.org/)^78^. Relevant pathways that contain five or more genes were selected. For data integration, total protein counts were fitted to a gaussian kernel to obtain expressed proteins^79^.

### Data integration

#### Input data for network generation

To reconstruct the intracellular signaling in PYK2-KD and FAK-KD tumor cells, two *a priori* interactions datasets were used. All possible signaling interactions known to occur in humans were obtained from the core protein-protein interaction (PPI) layer of the OmniPath collection using the ‘OmnipathR’ R package^80,81^. Only directed and signed interactions were included. Subsequently, upstream interactions between FAK or PYK2 and their interacting human binding proteins were identified by filtering for interactions stemming from FAK and PYK2. Interactions between human transcription factors (TFs) and their target genes (TG) were obtained from the DoRothEA collection using the DoRothEA R package^82,83^. Only signed interactions of the top three confidence levels (A, B, C) were included.

#### Causal network construction and visualization

The ViralLink pipeline^84^ was adapted to construct causal networks representing the intracellular signaling in PYK2-KD and FAK-KD tumor cells. Briefly, the list of expressed proteins in the PYK2-KD and FAK-KD cells was used to filter the *a priori* molecular interactions from OmniPath and DoRothEA, to obtain PPI and TF-TG sub-networks where both interacting molecules are expressed (described as “contextualised” networks). Transcription factors regulating the differentially expressed genes in PYK2-KD or FAK-KD were predicted using the contextualised DoRothEA TF-TG interactions and scored as described in^84^. Separate networks were built to construct the signal in PYK2-KD in the presence of FAK (FAK network), or in FAK-KD in the presence of PYK2 (PYK2 network). To build the FAK causal network, human binding proteins of FAK obtained from OmniPath^80,81^ were connected to TFs regulating DEGs in PYK2-KD *vs.* control through the contextualised PPIs in PYK2-KD using the network diffusion approach Tied Diffusion Through Interacting Events (TieDIE)^85^. Similarly, to build the PYK2 to FAK causal network, human binding proteins of PYK2 obtained from OmniPath^80,81^ were connected to TFs regulating DEGs in FAK-KD *vs*. control through the contextualised PPIs in FAK-KD using TieDIE. The final reconstructed networks contain “nodes”, which refers to the interacting partners, and “edges”, which refers to the interaction between nodes. Nodes include FAK or PYK2 and their human binding proteins, intermediary signalling proteins, TFs and differentially expressed genes in PYK2-KD or FAK-KD, respectively. Edges include activatory or inhibitory interactions. Network visualisations of PYK2 and FAK causal networks were performed using Cytoscape 3.8.2^86,87^. To visualise differences and overlaps in intracellular signalling, the separate built networks were subsequently joined using the “merge” function within Cytoscape. Nodes and edges were annotated according to their involvement in FAK or PYK2 causal networks.

#### Network functional analysis

A network-based pathway enrichment analysis was carried out on the separate PYK2 and FAK networks using the ANUBIX algorithm^51^. Using this approach, human proteins of the reconstructed causal network are tested for enrichment against a set of ANUBIX-provided Reactome and KEGG pathways with contextualised PPI networks as the input. All functions with q value ≤ 0.05 are considered significantly overrepresented, and pathways that contain five or more proteins were selected. To investigate how specific altered signalling pathways driven by FAK in PYK2-KD or by PYK2 in FAK-KD affect downstream functions of differentially expressed genes (DEGs), an in-built R script was created to construct function-specific subnetworks.

#### Invadopodia and focal adhesion analyses

To investigate the extent by which specific altered signaling pathways in PYK2 to FAK and FAK to PYK2 networks contribute to invadopodia and focal adhesions, the enrichment of invadopodia or focal adhesion-related proteins in PYK2 and FAK networks was assessed by performing an unranked hypergeometric test using the R package hypeR^88^. For this analysis, a list of manually curated invadopodia genes was generated by literature mining using GLAD4U, GeneShot, and ALS (Cytoscape)^89,90^. For generating the focal adhesions list, GLAD4U, GeneShot, ALS and the adhesome database^91^ were used. Each list was tested against the list of nodes in the FAK and PYK2 casual networks using the contextualised PPI networks as background and a False Discovery Rate (FDR) cut-off of 0.01. Subsequently, invadopodia and focal adhesion specific sub-networks were created as described above to visualise how specific altered signaling pathways driven by FAK in PYK2-KD or by PYK2 in FAK-KD affect downstream invadopodia or focal adhesion-related functions.

#### Statistical analysis

Statistical analysis was performed using GraphPad Prism 8.0. For all experiments, statistical significance was calculated using one-way ANOVA followed by Dunnett’s posthoc test. For patient database analysis, statistical significance was calculated using log-rank and Student’s *t*-test implemented in MATLAB. Values were considered significant if P<0.05. For all figures, *P<0.05, **P<0.01, ***P<0.001. Error bars represent standard errors of the mean (SEM).

