## Supplementary Figures for "FAK family proteins regulate *in vivo* breast cancer metastasis *via* distinct mechanisms"

**SUPPLEMENTARY MATERIAL**

**Supplementary Figure 1.** Enrichment analysis of transcriptome in PYK2-KD2 and FAK-KD1 tumors, that was measured by NanoString. Shown is the expression of genes included in cell cycle (A), mTOR signaling (B), MET signaling (C), apoptosis (D), and autophagy (E).

**Supplementary Figure 2.** Cartoon description of the data integration strategy. Details of the integration appear in the Results and in Materials and Methods sections.

**Supplementary Figure 3. PYK2 causal network.** PYK2 causal signaling network, demonstrating the role of PYK2 in intracellular signaling of breast tumor cells, in absence of FAK. The network contains three layers: PYK2 layer (PYK2 and its direct protein interactors), protein-protein interaction layer, and transcription factors and their differentially expressed genes layer. The shape of nodes indicates their association with a specific layer; direction of the interaction is indicated by the arrow shape (delta shape for activation/T shape for inhibition); the color of nodes indicates the difference in protein abundance, and the size of nodes indicates Log2 mean protein count.

**Supplementary Figure 4. FAK causal network.** FAK causal signaling network, demonstrating the role of FAK in breast tumor cell signaling in absence of PYK2. The network contains three layers: PYK2 layer (PYK2 and its direct protein interactors), protein-protein interaction layer, and transcription factors and their differentially expressed genes layer. The shape of nodes indicates their association with a specific layer; direction of the interaction is indicated by the arrow shape (delta shape for activation/T shape for inhibition); the color of nodes indicates the difference in protein abundance, and the size of nodes indicates Log2 mean protein count.

**Supplementary Video 1.** Representative field of a tumor cell with invadopod-like protrusion. Green, tumor cells; red, blood vessels; blue, collagen fibers. Scale bar, 10 μm.

**Supplementary Video 2.** Representative field of tumor cells exhibiting a fast-locomotion phenotype, in a region far from blood vessels. Green, tumor cells; red, blood vessels; blue, collagen fibers. The video relates to Figure 7A. Scale bar, 50 μm.

**Supplementary Video 3.** Representative field of a tumor cell exhibiting a fast-locomotion phenotype, in a region far from blood vessels. Green, tumor cells; red, blood vessels; blue, collagen fibers. Scale bar, 25 μm.

**Supplementary Video 4.** Representative field of tumor cells exhibiting a slow-locomotion phenotype, in a region close to a blood vessel. Green, tumor cells; red, blood vessels; blue, collagen fibers. The video relates to Figure 7B. Scale bar, 50 µm.

**Supplementary Video 5.** Representative field of a tumor cell exhibiting a slow-locomotion phenotype, in a region close to a blood vessel. Green, tumor cells; red, blood vessels; blue, collagen fibers. Scale bar, 25 µm.

**Supplementary Table 1.** Differentially expressed genes and proteins from PYK2-KD *vs.* control and FAK-KD *vs.* control xenograft mammary tumors.

**Supplementary Table 2.** RNA sequencing and mass spectrometry pathway enrichment analyses. Pathways containing 5 genes/proteins or above are presented.

**Supplementary Table 3.** PYK2/FAK network nodes and enrichment analysis of PYK2/FAK network proteins. Pathways containing 5 proteins or more are presented.

**Supplementary Table 4**. Curated invadopodia and focal adhesion genes and their relevant nodes from Pyk2 and FAK networks. Invadopodia genes were curated from GLAD4U, ALS, GeneShot, and from Hoshino *et al.*[1], and Hastie and Sherwood[2]. Focal adhesion genes were curated from GLAD4U, ALS, GeneShot, and from Zaidel-Bar *et al.*[3].

**Supplementary Table 5.** Summary of selected PYK2/FAK-associated *in vivo* signaling pathways mentioned in the paper.

**Supplementary Figure 1**

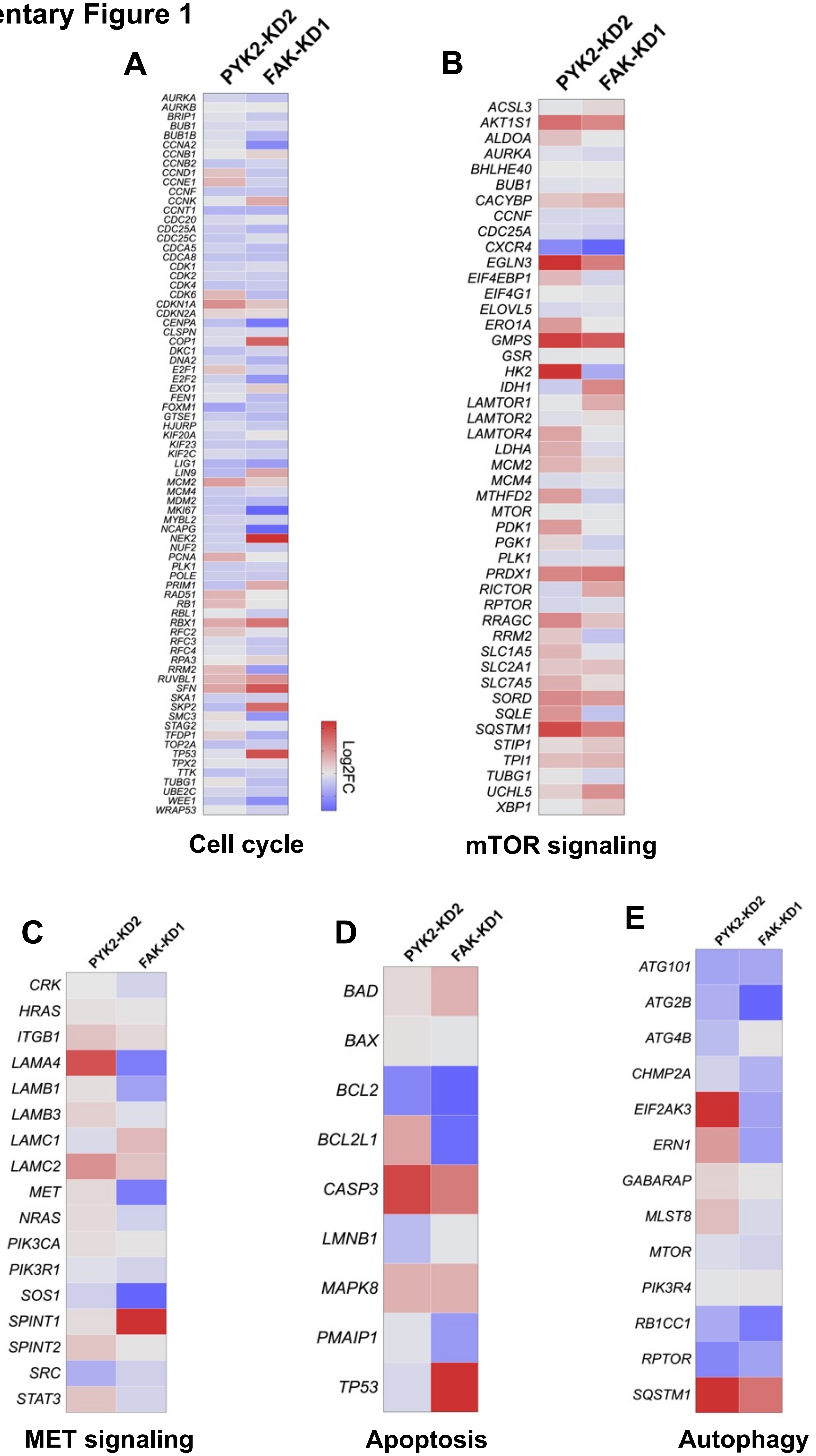

### Supplementary Figure 2

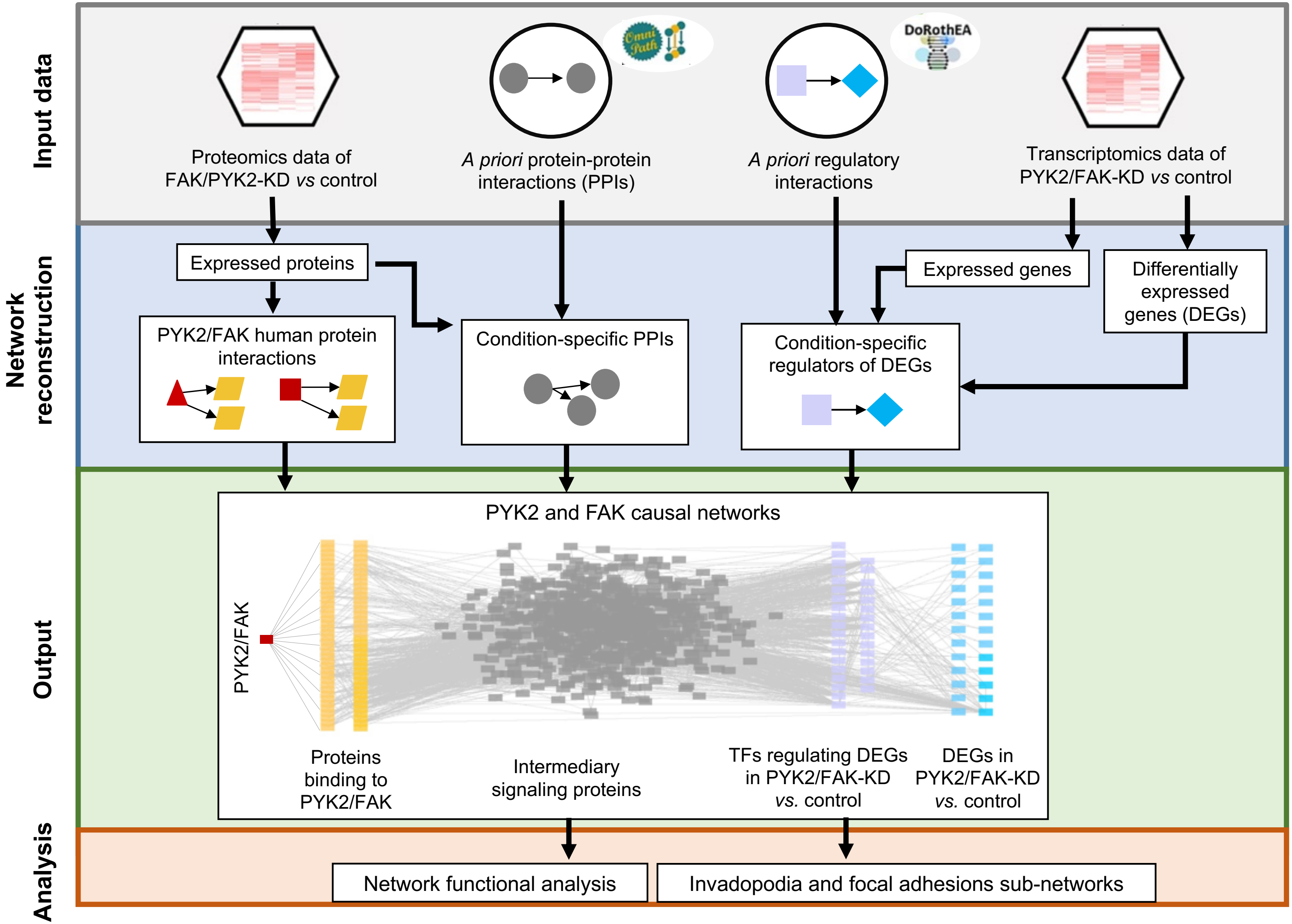

**Supplementary Figure 3**

#### PYK2 network

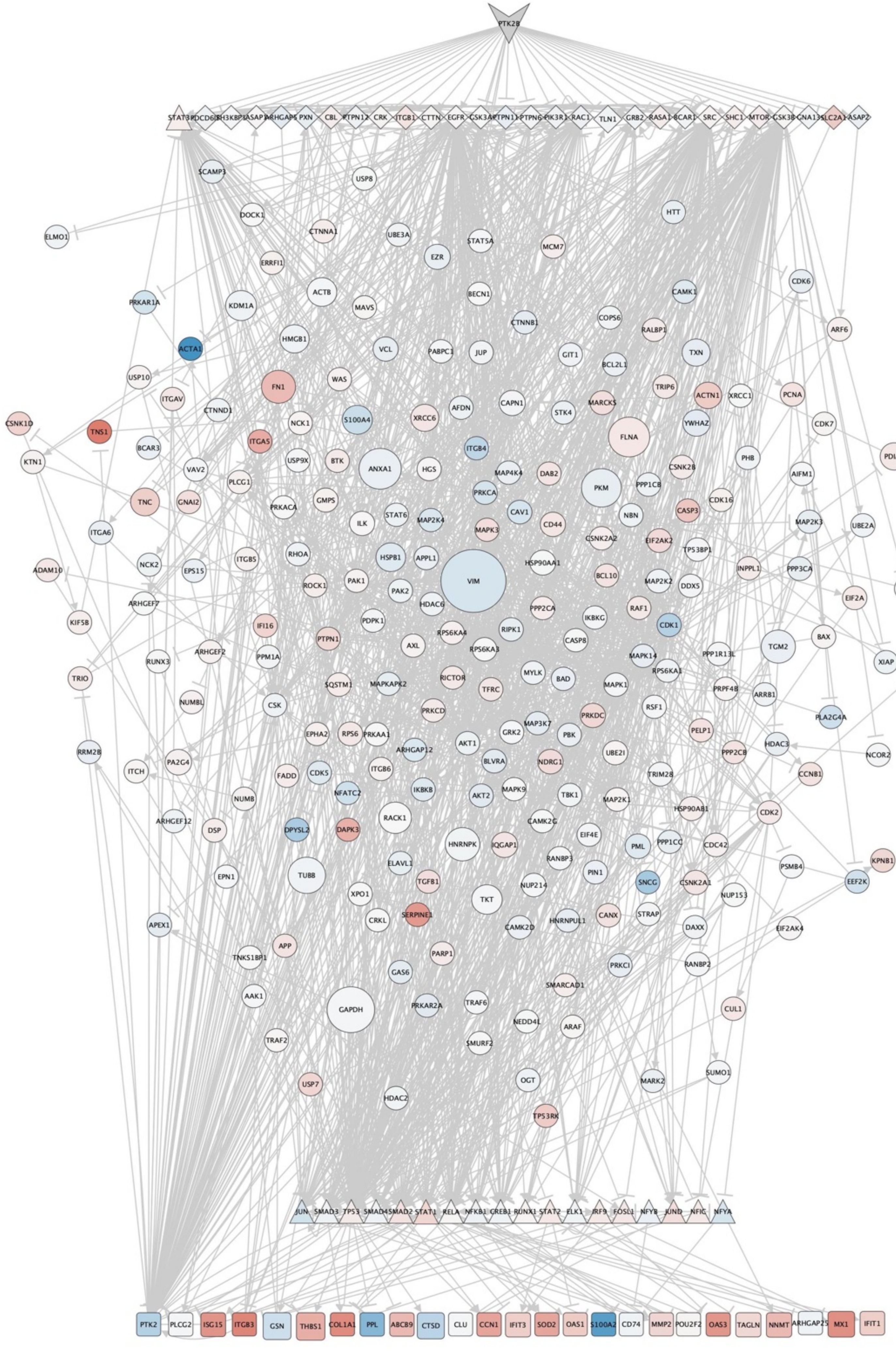

**Supplementary Figure 4**

#### FAK network

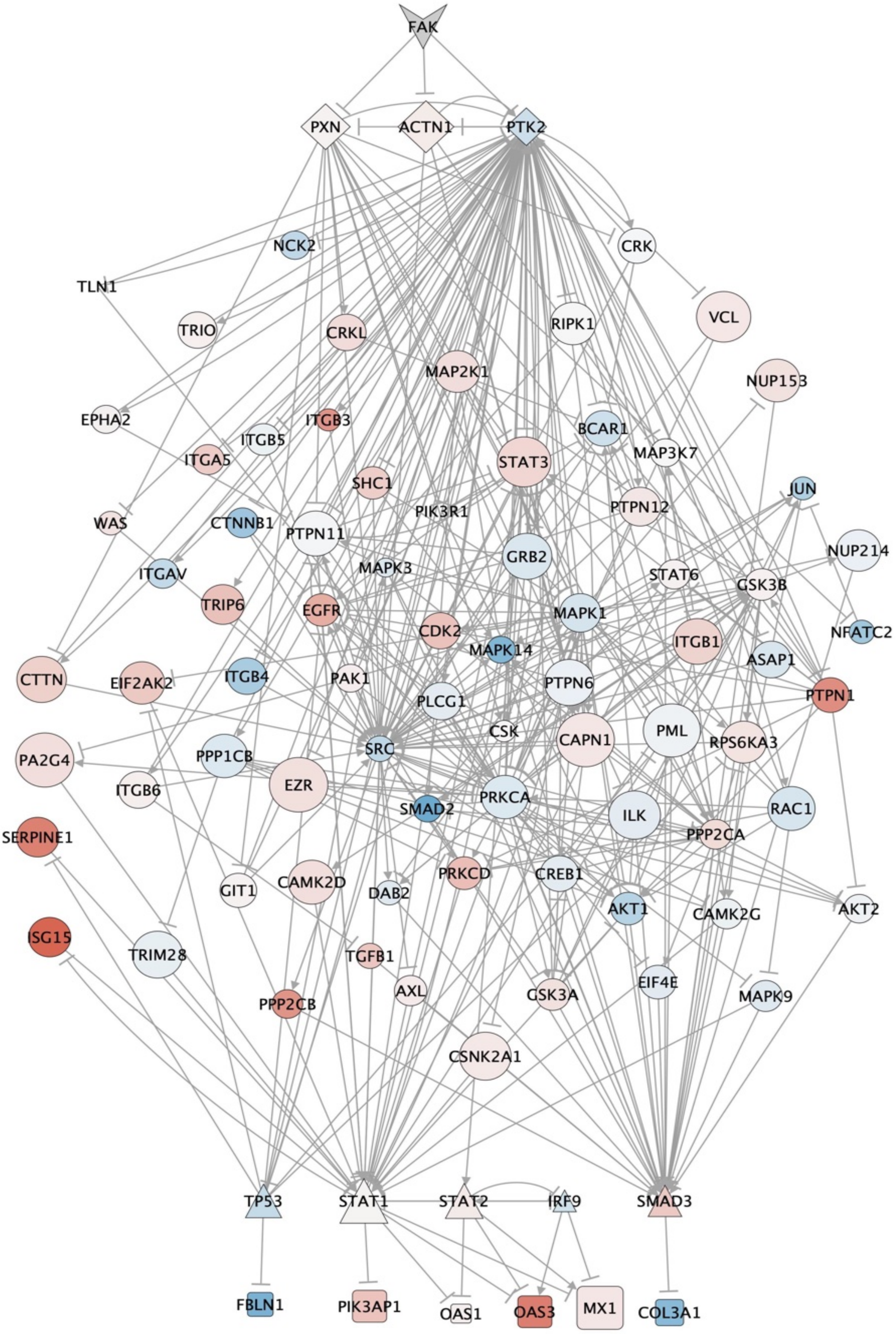
